# The miR319-targeted TCP transcription factors play crucial roles in the establishment of Arabidopsis thaliana shoot architecture in response to carbon and nitrogen availability

**DOI:** 10.64898/2026.09.08.750043

**Authors:** François Barbier, Isabella Amor, The Dan Pham, Thibaut Perez, Bin Luo, Liping Tang, Alessandro Silvestri, Ignacio Rubio Somoza, Michael Nicolas, Pilar Cubas, John Paul Alvarez, Franziska Fichtner, Christine Beveridge

## Abstract

Shoot branching is a highly plastic developmental process that allows plants to adjust their architecture to environmental conditions, such as carbon and nitrogen availability. Although the class II TCP transcription factor BRANCHED1 is a well-established repressor of shoot branching, the contribution of other TCP factors during this process remains less well understood. Here, we show that the miR319-targeted TCP module promotes shoot branching in *Arabidopsis thaliana*. Overexpression of *miR319* reduced rosette branch production, whereas repression of TCP3 activity or triple knockout (KO) mutation in *TCP3*, *TCP4* and *TCP10* inhibited branching. Conversely, expression of a miR319-resistant *TCP3* triggered a highly branched phenotype, indicating that TCP3 and related miR319 targets act as positive regulators of shoot branching. Genetic analysis with the strigolactone-deficient *max4* mutant showed that strigolactones mediate, at least partly, the reduced branching observed in the miR319 overexpressing line and in the *tcp3,4,10* triple mutant. Transcriptomic and DNA-binding analyses indicated indirect effect of TCP3 on strigolactone biosynthesis genes. Instead, *miR319* overexpression increased the sensitivity of branching to nitrogen limitation and reduced nitrate uptake, thereby increasing the expression of strigolactone synthesis genes. We further show that carbon starvation reduced the transcription levels of miR319-targeted *TCPs*, independently of miR319 accumulation, and that TCP3 may connect carbon availability to sugar signalling via HEXOKINASE1. Together, our results identify the miR319-targeted TCP module as a positive regulator of shoot branching that links plant architecture to carbon and nitrogen availability.

## Introduction

Shoot branching is a highly plastic trait that plays a crucial role in shaping shoot architecture in response to the environment and is therefore critical for plant fitness and crop yield. For example, shoot branching is particularly sensitive to carbon and nitrogen availability. Conditions that increase carbon availability, such as high light and elevated CO₂, generally favour shoot branching, whereas low light or nitrogen limitation inhibits this process (de Jong et al., 2014; Zhou et al., 2021; Fichtner et al., 2022; Sakioka and Yoneyama, 2025). Understanding how carbon and nitrogen availability regulate shoot branching is therefore critical for determining how plants adjust their architecture to adapt to the environment.

Shoot branching in flowering plants is mainly driven by the outgrowth of axillary buds into branches. Axillary buds are compact structures produced at the axils of leaves and contain a meristem and leaf primordia. After their formation, axillary buds are maintained in a quiescent or dormant state. While their formation is genetically determined, the release of axillary buds from dormancy is tightly regulated by networks of molecular and physiological signals that are controlled by environmental cues (Barbier et al., 2019b; Beveridge et al., 2023).

In herbaceous plants, one of the most important mechanisms controlling axillary bud dormancy is apical dominance, a phenomenon through which the growing shoot apex inhibits the outgrowth of axillary buds along the stem (Barbier et al., 2017; Luo et al., 2021; Beveridge et al., 2023). Apical dominance is maintained through two systemic signals: auxin and sugars. On one hand, the growing shoot tip produces a flow of auxin that is transported basipetally in the stem and inhibits bud outgrowth (Wang et al., 2018; Barbier et al., 2019b; Luo et al., 2021). On the other hand, the growth of the shoot apex generates a strong sink that limits sugar availability to axillary buds, thereby maintaining their dormancy (Mason et al., 2014; Barbier et al., 2015b; Beveridge et al., 2023). When the plant is decapitated, auxin is progressively depleted from the stem and sugar fluxes are redirected toward axillary buds, triggering their outgrowth (Mason et al., 2014; Fichtner et al., 2017; Cao et al., 2023).

Auxin produced by the growing apex and flowing in the main stem does not enter axillary buds and therefore acts indirectly to inhibit their outgrowth (Hall and Hillman, 1975). Instead, auxin represses the synthesis of cytokinins, a class of hormones that induce bud outgrowth (Tanaka et al., 2006; Barbier et al., 2019b), and induces the synthesis of strigolactones (SL), a class of hormones that repress bud outgrowth (Gomez-Roldan et al., 2008; Brewer et al., 2009). Conversely, sugars have also been shown to promote cytokinin accumulation and repress SL perception to promote axillary bud outgrowth (Barbier et al., 2015a; Bertheloot et al., 2020; Barbier et al., 2021; Salam et al., 2021; Patil et al., 2022). Cytokinins and SL control bud outgrowth via different downstream mechanisms, including the regulation of different transcription factors and the induction of auxin export from axillary buds, which promotes sustained bud outgrowth (Bennett et al., 2006; Waldie and Leyser, 2018; Cao et al., 2023; Nahas et al., 2024).

In addition to their trophic roles, sugars control plant metabolism and development via a network involving different signalling components (Fichtner et al., 2021b; Li et al., 2021). Sugars have been reported to induce shoot branching, at least partly, by inducing accumulation of the sucrose-specific signalling metabolite trehalose 6-phosphate (Tre6P) in axillary buds (Fichtner et al., 2017; Fichtner et al., 2021a). A glucose-specific pathway mediated by HEXOKINASE1, an enzyme that acts as a glucose sensor in plants (Moore et al., 2003; Cho et al., 2006) was also shown to be positively involved in the control of shoot branching in *Arabidopsis thaliana* (Barbier et al., 2021).

Transcription factors play important roles in the integration of multiple signals that control axillary bud outgrowth, most notably the class II TEOSINTE BRANCHED1/CYCLOIDEA/PCF (TCP) transcription factor BRANCHED 1 (BRC1) (Aguilar-Martínez *et al*., 2007) and its monocot ortholog TEOSINTE BRANCHED1 (TB1) (Doebley et al., 1997; Wang et al., 2019a). *BRC1* is strongly expressed in dormant axillary buds and have been demonstrated to inhibit bud outgrowth in many flowering plant species (Aguilar-Martínez et al., 2007; Wang et al., 2019a). The expression of this transcription factor is repressed by signals promoting bud outgrowth, such as sugars and cytokinins, and is induced by signals repressing this process, such as SL and low red:far-red light ratio, placing BRC1 in a central position in the integration of signals controlling shoot branching (Barbier et al., 2019b; Wang et al., 2019b). The importance of BRC1 in shoot branching and tillering makes this transcription factor a key target for crop improvement (Doebley et al., 2006).

However, BRC1 is not the only TCP transcription factor to control shoot branching. In Arabidopsis, BRC2 (TCP12), the close paralog of BRC1 (TCP18), also plays an inhibitory role ins shoot branching, although quite minor compared to BRC1 (Aguilar-Martínez et al., 2007). In the same species, the class I TCPs TCP14 and TCP15 were recently reported to promote shoot branching in *Arabidopsis thaliana*, notably in response to light quality (Gastaldi et al., 2024). A homologue of TCP14 in apple tree was also recently shown to promote bud break in response to chilling temperatures (Zhang et al., 2026). The miR319-targeted class II TCPs (*TCP2, TCP3, TCP4, TCP10* and *TCP24*) also seem to have a role in the regulation of shoot architecture. These TCPs are divided into two clades: one including TCP2 and TCP24, and one including TCP3, TCP4 and TCP10 (Martín-Trillo and Cubas, 2010). It was recently reported that the *tcp3,4,10* triple KO mutant displays a decreased shoot branching phenotype, showing that this other clade of miR319-targeted TCPs is also involved in the control of shoot branching (Huang et al., 2026). Furthermore, TCP4 was shown to interact physically with SMXL6 (Huang et al., 2026), a repressor of SL signalling (Wang et al., 2015), to directly repress *BRC1* expression, indicating crosstalk between TCP4 and the SL signalling pathway. However, overexpressing *TCP4* is not sufficient to induce shoot branching in wild-type plants but can slightly increase shoot branching in SL mutants. In contrast, overexpressing its closest homologue *TCP3*, triggers an extremely bushy phenotype in arabidopsis (Li and Zachgo, 2013), suggesting that TCP3 and TCP4 play different roles in shoot branching. In rice, the homologue of AtTCP2 was reported to promote tillering, while miR319 inhibits this developmental process (Wang et al., 2021), which was also reported for bentgrass (*Agrostis stolonifera*) and switchgrass (*Panicum virgatum*) (Zhou et al., 2013; Liu et al., 2020). In *Camellia sinensis*, the close homologue of AtTCP2 was suggested to be involved in the regulation of bud dormancy based on the positive correlation between its expression and bud dormancy release, which was concomitant with a decrease in the expression of *miR319*. Together, these observations suggest that BRC1 is not the only TCP transcription factor to control shoot branching and that the TCP regulatory network controlling this process is broader than previously thought.

Despite these observations, the precise position of miR319 and its targets in the shoot branching network remains poorly understood, in particular in relation to sugar availability and SL signalling, two central regulators of bud outgrowth (Bertheloot et al., 2020; Patil et al., 2022; Barbier et al., 2023; Fichtner et al., 2024). Here, we focused more specifically on TCP3, whose overexpression was previously reported to trigger a strong bushy phenotype in *Arabidopsis thaliana* (Li and Zachgo, 2013). Using a combination of genetic and physiological approaches, we investigated the role of miR319, TCP3 and its close homologues in the control of shoot branching in the context of nutrition. Our results provide evidence that miR319 targets act as positive regulators of shoot branching, are specifically induced by carbon availability and mediate the response to nitrogen status, thereby revealing a previously underappreciated role for these transcription factors in the regulation of plant architecture in response to nutrients.

## Material and Methods

### Plant material and growth conditions

All experiments were performed on *Arabidopsis thaliana* Columbia-0 (Col-0) or Landsberg *erecta* (L*er*) ecotypes. The miR319 promoter-GUS reporter lines were previously published (Nag et al., 2009). The *jaw-D* line was previously described and carries a 35S constitutive promoter inserted upstream of the *miR319a* gene (Palatnik et al., 2003). The *35S:miR319a* line in the L*er* background was previously described (Alvarez et al., 2016). The *35S:TCP3-SRDX* and *35S:mTCP3* lines were previously published (Li and Zachgo, 2013). The triple *tcp3,4,10* mutant was obtained from Tomotsugu Koyama and previously published (Koyama et al., 2010). The *proSTM>>GUS* and *proSTM>>miR319* transactivation lines were obtained by crossing the previously published *proSTM:LhG4* line to the *proOP:GUS* and *proOP:miR319* (Moore et al., 1998; Efroni et al., 2008; Yadav et al., 2010). The SL-deficient *max4-1* mutant was previously described (Sorefan et al., 2003).

Plants were grown in growth chambers or cabinets with a light intensity between 120 and 150 µmol photons m⁻² s⁻¹ PPFD, a 16 hr photoperiod, and a temperature between 18 and 21°C at night, and between 21 and 23°C during the day. In most experiments, plants were grown using UQ23 substrate, which consists in a mixture of fine pine bark, coco peat, sand and vermiculite. In figure 5 and 7, plants were grown using Jiffy PP4 substrate, which consists in a mixture of sphagnum peat, clay and coco fibre supplemented with 50% of sand.

### Generation of the brc1 CRISPR knockout line

After a first round of experiments, using a cross between *jaw-D* and *brc1-2*, we noticed that the T-DNA of the *brc1-2* allele caused silencing of the *miR319a* overexpression. To avoid this effect, we generated a CRISPR knockout (KO) allele of *brc1*. For this, we used an intronized Cas9 (Grützner et al., 2021), under the control of the *RPS5a* promoter. The vector’s cassette contained a cloning site for a single guide RNA and an *OLEOSINp:RFP* cassette for seed selection (Ursache et al., 2021). The gRNA was designed using the webtool Breaking-Cas (https://bioinfogp.cnb.csic.es/tools/breakingcas, Oliveros et al., 2016). T0, RFP-positive seeds were selected under the stereoscope and genotyped by PCR using primers listed in Supp. Tables S1. In mutant T2, RFP-negative, Cas9-free seeds were selected. The *brc1-8* CRISPR mutant has an A insertion which causes a frameshift and an early stop codon, leading to a CDS predicted to encode a truncated protein of 88 aa, lacking the TCP and R domains (Supp. Fig. S1). To reduce the number of potential off-target mutations the mutant was backcrossed once with WT Col-0.

### Creation of the proBRC1:TCP3-SRDX line

For the expression of *TCP3-SRDX* under the *BRC1* promoter (1kb upstream the TSS) the *pGreenII pBRC1:GUS* plasmid was used (as described in Fichtner et al., 2021). The *TCP3* open reading frame without introns was amplified from *Arabidopsis thaliana* Col-0 cDNA and the ERF-associated amphiphilic repression (EAR) motif repression domain (SRDX) was added during the PCR amplification by primer extension. All primer sequences are given in Supplementary Table 1. The *GUS* sequence was then replaced by the *TCP3-SRDX* open reading frame using the NheI restriction sites of the *TCP3-SRDX* PCR product added during amplification and the compatible SpeI sites in the *pGreenII pBRC1:GUS* plasmid. The *pGreenII pBRC1:TCP3-SRDX* construct was introduced into *Arabidopsis thaliana* Col-0 by *Agrobacterium tumefaciens*-mediated transformation using strain GV3101(pMP90) (Koncz and Schell, 1986) and the floral dip standard protocol (Clough and Bent, 1998). Primary transformants were selected by growing seedlings under long day conditions on half-strength MS medium containing 10 mg/L phosphinothricin. Lines that showed a 3:1 segregation of resistant:susceptible individuals in the T_2_ generation, indicating a single transgenic locus, were chosen for further propagation. Progeny from such lines were screened in the T_3_ generation to identify homozygous lines for each transgene by survival assays.

### Phenotypic characterisation

Shoot branching was determined by scoring the number of primary rosette branches longer than 1 cm produced every two days after bolting, the stage at which shoot branching usually starts in *Arabidopsis thaliana*. In figure 2D, the number of primary, secondary, tertiary, quaternary and quinary rosette and cauline branches was scored when the plants were starting to senesce.

### GUS staining

GUS staining was performed by collecting samples directly in staining buffer containing 0.5 mM X-Gluc dissolved in DMSO. The staining buffer consisted of 68 mM Na_2_HPO_4_, 32 mM NaH_2_PO_4_, 0.5 mM potassium ferrocyanide, 0.5 mM potassium ferricyanide and 0.2% Triton X-100. Samples were vacuum infiltrated for 10 to 15 min, and then incubated at 37°C for up to 8 hrs. Staining was stopped by rinsing samples twice with water. Samples were then cleared in 70% ethanol with gentle agitation before imaging.

### RNA extraction and gene expression analyses by Real-Time Quantitative PCR (RT-qPCR)

Total RNA was extracted from frozen-ground samples or from freeze-dried ground samples using a phenol/chloroform-free CTAB-based method as previously published (Barbier et al., 2019a) or by using the NucleoSpin RNA extraction kit (Macherey-Nagel, Germany). For gene expression analysis using RT-qPCR, diluted cDNA was then used as a template for quantitative real-time PCR following the manufacturer’s instructions (SensiFAST™ SYBR® No-ROX, Meridian Bioscience) or (TB Green® Premix Ex Taq^TM^, Takara). Product amplification was monitored with a CFX384 TouchTM RealTime PCR Detection System (Bio-Rad) or LightCycler® 384 (Roche). *TUBULIN3* and *BetaACTIN* were used for normalisation. Primers used in this study are listed in Supp. Table S1.

### Small RNA sequencing

Libraries were prepared from the same RNA samples and sequenced on NovaSeq 6000 generating 75 bp paired end reads with a target depth of 25 million reads per sample. Small RNA sequencing reads were quality-checked using FastQC (v0.11.5) and adapter/quality trimmed with Trim Galore (v0.4.5) using the parameters “-q 20 --length 16 --max_length 33”, which also generated post-trim FastQC reports. Trimmed reads were mapped to small RNA–generating loci annotated in the Plant Small RNA Genes database (https://plantsmallrnagenes.science.psu.edu/genomes.php?genomes_id=49), which provides consistent annotation of sRNA loci across plant species. Read alignment was performed with HISAT2 (v2.1.0) using a single-end alignment strategy and allowing up to 100 reportable alignments per read (-k 100), followed by filtering to retain reads with zero mismatches and subsequent sorting using SAMtools (v1.15.1). Locus-level read quantification was carried out with HTSeq (v0.12.4) using htseq-count with settings “-s no-t exon -- nonunique all-a 0 --secondary-alignments score”. Differential expression analysis of sRNA loci was performed using DESeq2 with default parameters. Small RNA loci with padj < 0.05 were considered differentially expressed. Annotation information including sRNA sequence, family classification (miRNA, siRNA, or other), and overlapping gene features was obtained from the Plant Small RNA Genes database.

### Analysis of TCP3 DAP-seq data

To assess the binding of TCP3 on different genomic loci, publicly available DNA affinity purification sequencing (DAP-seq) data were obtained from the Plant Cistrome Database (https://neomorph.salk.edu/dap_web/pages/index.php) (O’Malley et al., 2016). TCP3 binding tracks were visualised using the genome browser available on the database website. Candidate genomic loci were inspected manually for the presence of TCP3 DAP-seq peaks in the promoter and gene body regions as indicated in the figure legends. Browser screenshots of the relevant genomic regions were used to illustrate TCP3 binding profiles in the corresponding figures.

### Nitrate root uptake experiment

To assess nitrate root uptake capacity, arabidopsis seedlings were grown for 10 days on half-strength MS medium containing 1 mM ammonium nitrate, under the same growth conditions as described above. Seedlings were then incubated for 1 min in 0.1 mM CaSO₄ buffer before being transferred for 5 min to half-strength nitrogen-free MS medium containing 200 µM K¹⁵NO₃. Seedlings were then rinsed for 1 min in 0.1 mM CaSO₄ buffer. For each genotype, three roots were pooled together and dried before analysis. Isotopic enrichment in ¹⁵N was measured by EA-IRMS (Elementar, UK). Nitrate uptake was determined by normalising ¹⁵N enrichment to sample dry weight and to the duration of the influx assay.

### Statistical analyses

Significant differences between samples were determined using Student’s t-test or one-way analysis of variance (ANOVA) when appropriate, with α = 0.05. Significant differences are represented by asterisks for Student’s t-test or by letters for one-way ANOVA, as indicated in the figure legends. Correlations between gene expression values were assessed using Pearson correlation coefficients. For small RNA sequencing, differential expression analysis was performed using DESeq2 with default parameters, and small RNA loci with an adjusted P-value < 0.05 were considered differentially expressed. In all figures, data are given as means ± standard error or the mean (SE).

## Results

### miR319 inhibits shoot branching in arabidopsis

To investigate the role of miR319 in arabidopsis shoot branching, we first examined whether the three genes encoding for miR319 (*miR319a*, *miR319b* and *miR319c*) are expressed in tissues relevant to this developmental process using promoter-*GUS* fusion lines previously published (Nag et al., 2009). In contrast to observations in monocotyledonous plants where the expression pattern was ubiquitous (Wang et al., 2021), the expression pattern of the genes encoding for *miR319* in *A. thaliana* is extremely restricted as previously shown (Nag et al., 2009). In addition, close inspection of the arabidopsis rosette core shows that *miR319b* and *miR319c* are expressed in axillary buds (Fig. 1A), suggesting a role of these two miRNAs in axillary bud outgrowth and shoot branching. MiR319a was only detected in the stipular region at the base of young leaf primordia (Supp. Fig. 2A), making it less likely to be involved in the control of shoot branching.

**Figure 1.**
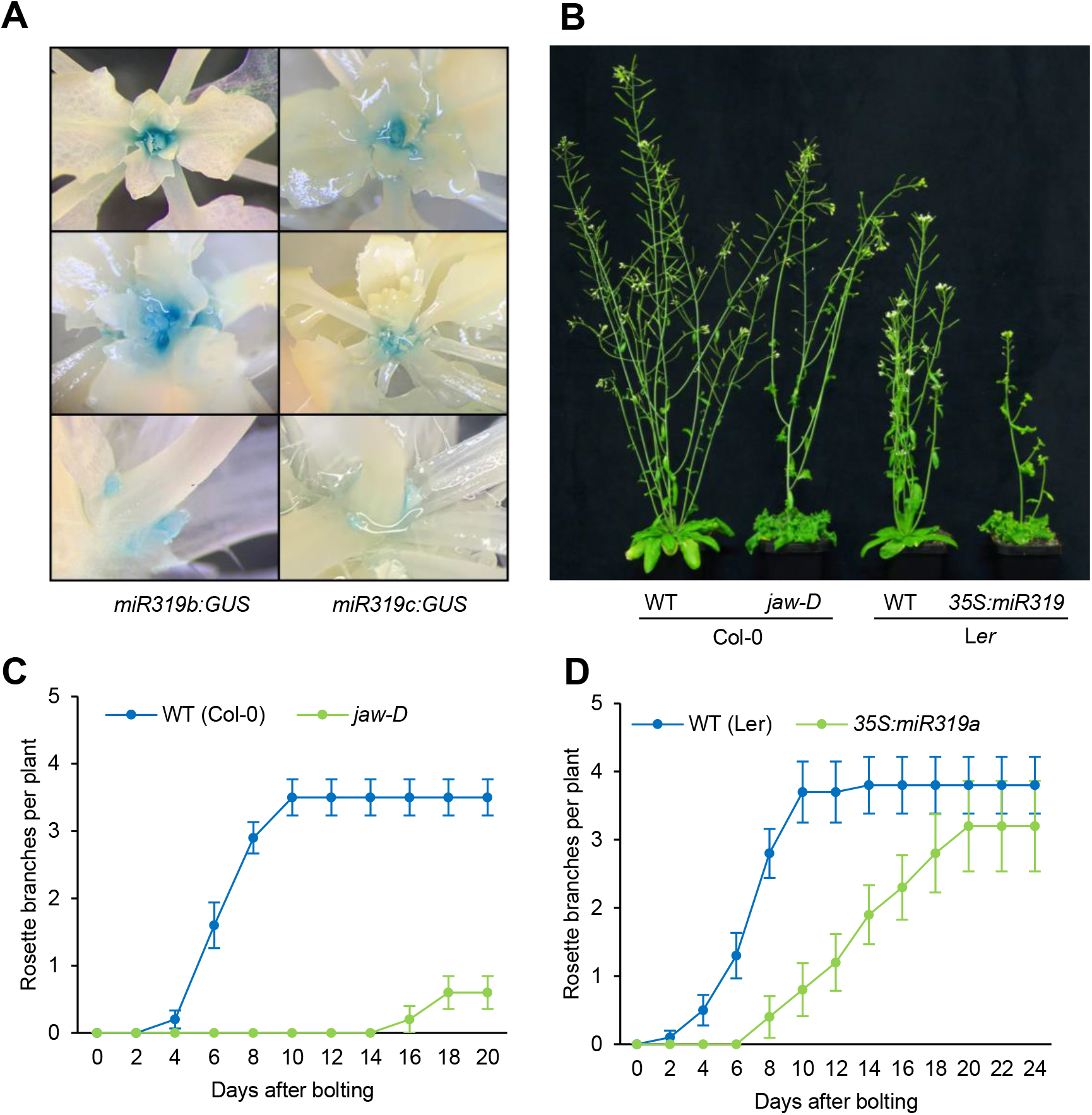
miR319 inhibits shoot branching in *Arabidopsis thaliana*. (**A**) GUS staining showing the expression pattern of *miR319b* and *miR319c* in the core of the *Arabidopsis thaliana* rosette at different stages of plant development. (**B**) shoot phenotype of ∼6 week-old plants over-expressing *miR319a* in Columbia-0 (Col-0; *jaw-D*) and Landsberg *erecta* (L*er; 35S:miR319*). (**C**) and (**D**) number of primary rosette branches emerging from Col-0 (C) and L*er* (D) plants over-expressing *miR319a*. Data are mean +/-S.E. (n=8-12 plants).

**Figure 2.**
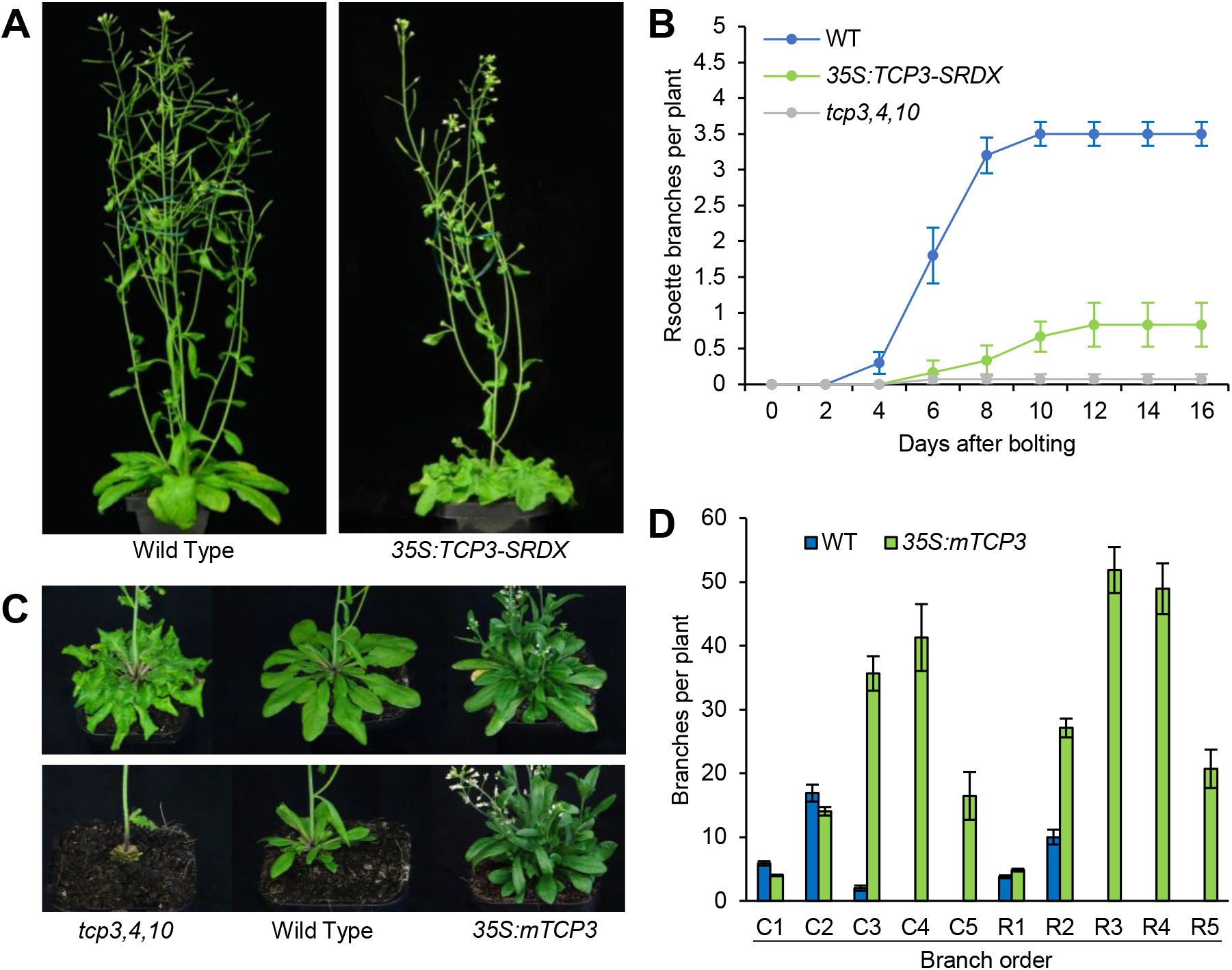
TCP3, TCP4 and TCP10 are positively involved in shoot branching in *Arabidopsis thaliana.* (**A**) shoot phenotype of Col-0 wild type and *35S:TCP3-SRDX* plants. (**B**) number of primary rosette branches emerging from wild-type Col-0 (WT), *35S:TCP3-SRDX* plants and *tcp3,4,10* triple mutant. Data are mean +/-S.E. (n=8-12 plants). (**C**) shoot phenotype of Col-0 wild type plant, *tcp3,4,10* triple KO mutant and *35S:mTCP3* expressing line, ∼7 days after bolting with (top picture) or without (bottom picture) their primary rosette leaves to show the secondary leaves emerging from the axillary buds. (**D**) final number of rosette (R) and cauline (C) branches of different orders in the WT Col-0 and the *35S:mTCP3* lines. Data are mean +/-S.E. (n=8 plants).

To test the role of miR319 in shoot branching, we characterised the shoot branching phenotype of lines overexpressing *miR319* in Col-0 and L*er* backgrounds, referred to as *jaw-D* and *35S:miR319*, respectively (Palatnik et al., 2003; Alvarez et al., 2016). Overexpression of *miR319* strongly reduces the production of primary rosette branches in both backgrounds (Fig. 1B-D). Rosette leaf removal also reveals that the growth of the secondary leaves produced by the axillary buds is inhibited in these lines (Supp. Fig. S2B), indicating that *miR319* overexpression not only inhibits the inflorescence outgrowth but also development of bud leaves, which happens first during branch emergence. These observations demonstrate that *miR319* inhibits arabidopsis bud development and branch outgrowth.

### The TCP targets of miR319 promote shoot branching

We then sought to characterise the role of the targets of *miR319* in arabidopsis shoot branching: *TCP2*, *TCP3*, *TCP4*, *TCP10*, and *TCP24*. To achieve this, we first tested whether overexpressing one of these targets fused to an SRDX motif that recruits TOPLESS and thereby represses transcription, could inhibit shoot branching. Using the *35S:TCP3-SRDX* line created by Li and Zachgo 2013 (Li and Zachgo, 2013), we observed a strong reduction in the number of rosette branches produced overtime in this line compared to the wild type (Fig. 2A-B). Given that ectopic overexpression of constitutive repressors could give rise to pleiotropic effects, we wanted to test whether knocking out several of the miR319 targets could also lead to an inhibition of shoot branching. To do so, we used a triple *tcp3,4,10* knockout (KO) line (Koyama et al., 2007). This line displayed a phenotype very similar to the *35S:TCP3-SRDX* line, with a strong inhibition of rosette branching (Fig. 2B-C).

To further test the involvement of the miR319 targets in shoot branching, we used a line that overexpresses a miR319-resistant version of TCP3, *35S:mTCP3* (Li and Zachgo, 2013). As previously reported, this line displayed very pleiotropic effects, including a strong increase of shoot branching. The secondary rosette leaves, characteristic of the first phase of rosette bud outgrowth, were much more visible compared to the wild type and the triple *tcp3, 4, 10* KO line for which the development of secondary leaves was strongly inhibited (Fig. 2C). Furthermore, we observed a strong increase in the order of branches produced, *i.e.* the number of successive branching levels that arose from the main axis (Supp. Fig. S3). Wild-type plants produced two orders of rosette branches and three orders of cauline branches, whereas *35S:mTCP3* produced up to five orders of branches for both cauline and rosette branches (Fig. 2D). Together, the increased shoot branching in the over-expressing line and the decreased branching in the KO mutant and suppressing line demonstrate that TCP3 and its paralogs are positively involved in shoot branching (Figure 2).

### Inhibition of TCP3 activity in axillary buds represses shoot branching

The observations in Figures 1 and 2 show that miR319 and its targets TCP3, TCP4 and TCP10 control shoot branching in arabidopsis. Because shoot branching plasticity is mainly driven by axillary bud outgrowth, we wanted to test whether suppressing TCP3 activity specifically in buds could repress shoot branching. To achieve this, we created transgenic lines in which the expression of the *TCP3-SRDX* construct (Fig. 2A-B) is driven by the *BRC1* promoter (Fig. 3A), as previously used (Fichtner et al., 2021a). Two independent *proBRC1:TCP3-SRDX* lines showed reduced rosette branch emergence, indicating that reduction of TCP3 activity in BRC1-expressing cells is sufficient to inhibit shoot branching.

**Figure 3.**
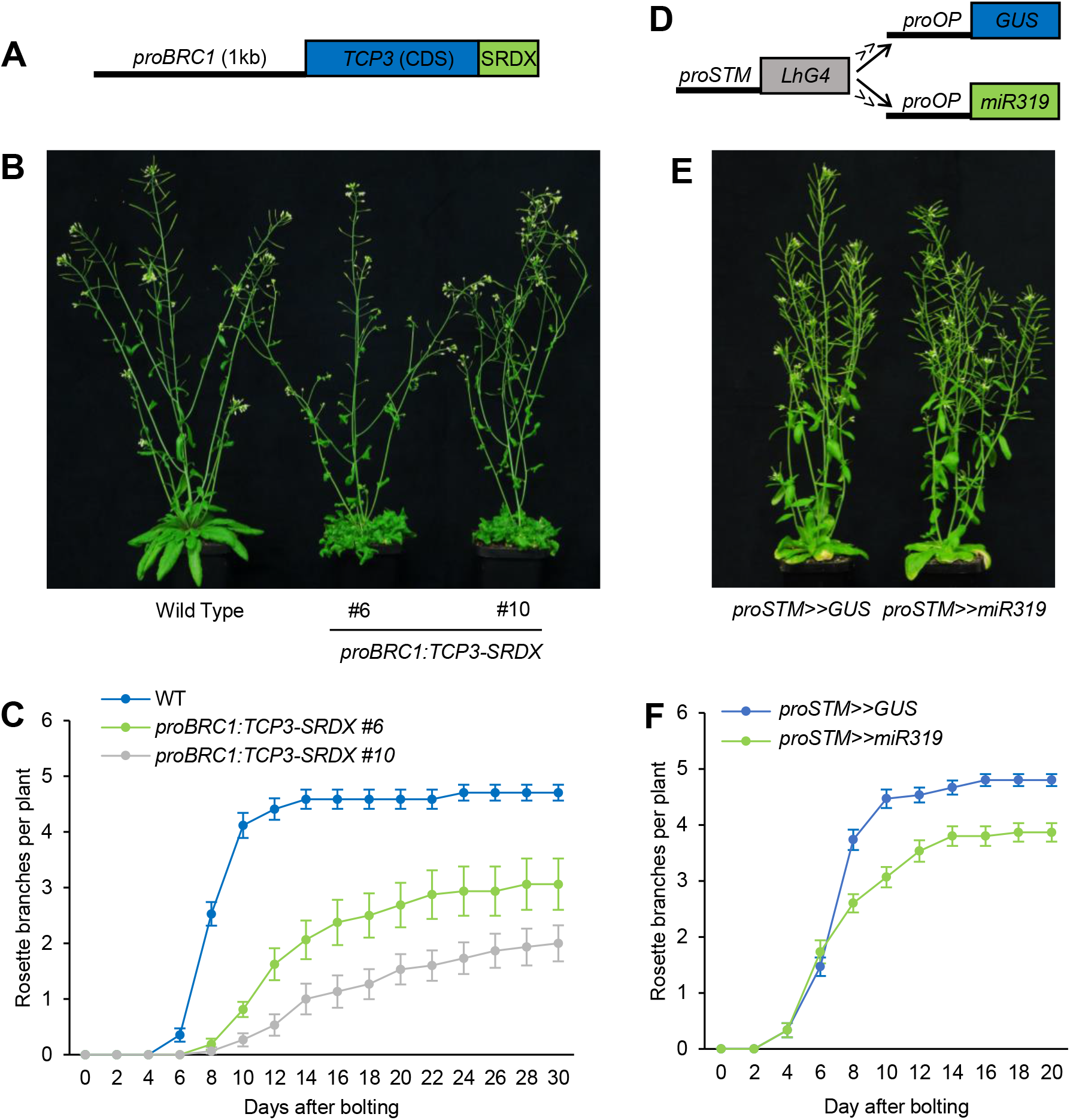
Localised expression of *TCP3-SRDX* and *miR319* inhibits shoot branching in *Arabidopsis thaliana*. (**A**) construct used to create the transgenic lines used in panel B and C. (**B**) Shoot phenotype of Col-0 wild type and two independent transgenic lines carrying the the *proBRC1:TCP3-SRDX* construct, two weeks after bolting. (**C**) number of primary rosette branches emerging from two independent line expressing the *proBRC1:TCP3-SRDX* construct and their wild-type Col-0. Data are mean +/-S.E. (n=8-12 plants). (**D**) transactivation system used in panels E and F. (**E**) shoot phenotype of the *proSTM>>GUS* and *proSTM>>miR319* plants. (**F**) number of primary rosette branches emerging from the *proSTM>>GUS* and *proSTM>>miR319* plants. Data are mean +/-S.E. (n=8-12 plants).

We also observed that both independent lines displayed crinkly waffled leaves reminiscent of the *35S:TCP3-SRDX* line (Fig. 2A), which is unexpected since the promoter used is not supposed to be expressed in leaves (Fichtner et al., 2021a). We suspected that the *TCP3-SRDX* may misregulate the expression of *BRC1*, thereby disturbing the expression pattern of the *proBRC1:TCP3-SRDX* transgene. To test this hypothesis we crossed the *proBRC1:TCP3-SRDX* #10 line to a line carrying the *proBRC1:GUS* construct. GUS staining analysis of *proBRC1:TCP3-SRDX* x *proBRC1:GUS* line confirmed that the *proBRC1:TCP3-SRDX* construct disturbs the spatial expression pattern of *BRC1*, leading to its expression in the base of the rosette leaves and more ectopic expression in very young leaves (Supp. Fig. S4A), likely explaining the leaf phenotype of the *proBRC1:TCP3-SR*DX lines.

Because the use of the *BRC1* promoter proved difficult for testing the local effect of the miR319 module in axillary buds (Fig. 3A and Supp. Fig. S4A), we used a transactivation approach to express *miR319a* under the control of *SHOOT MERISTEMLESS (STM)*, a gene specifically expressed in shoot meristems including axillary meristems but not in leaf primordia (Fig. 3D) (Long et al., 1996; Moore et al., 1998; Long and Barton, 2000; Efroni et al., 2008; Yadav et al., 2010). This *proSTM>>miR319* line displayed a more inhibited shoot branching phenotype than the *proSTM>>GUS* transactivation control (Fig. 3E-F), albeit without any effect on rosette leaf development (Supp. Fig. S4B). However, this inhibition was weaker compared to the observed effect of the *35S:miR319a* construct (Fig. 1), potentially due to the narrower expression pattern of *STM* (Long et al., 1996; Long and Barton, 2000) compared to the expression mediated by the *35S* promoter (Fig. 1A). Altogether, the data presented in Fig. 3 indicate that inducing *miR319* and repressing TCP3 activity in restricted regions including axillary buds can inhibit shoot branching.

### SL deficiency alleviates the inhibitory effect of miR319 on shoot branching

SLs are major endogenous repressors of bud outgrowth that act, in large part, by inducing the expression of the transcription factor *BRC1*, which as mentioned above, also belongs to the class II TCPs. We therefore wanted to test whether SLs repress shoot branching by regulating the expression of the miR319-targeted *TCP*s. To achieve this, we first quantified the expression of the three *miR319* precursors together with their *TCP* targets in the rosette of three-week-old *max4-1* mutant plants, which is deficient in SL. This analysis shows that the expression of these genes is similar between the *max4* mutant and its corresponding wild type Col-0 (Fig. 4A). The same observation was made when determining the expression of these genes in three-week-old axillary-bud enriched rosette cores of the L*er* ecotype with *max4-10* mutant (Supp. Fig. S5). These results clearly indicate that the SL signalling pathway does not control the expression of *miR319* or its targets.

**Figure 4.**
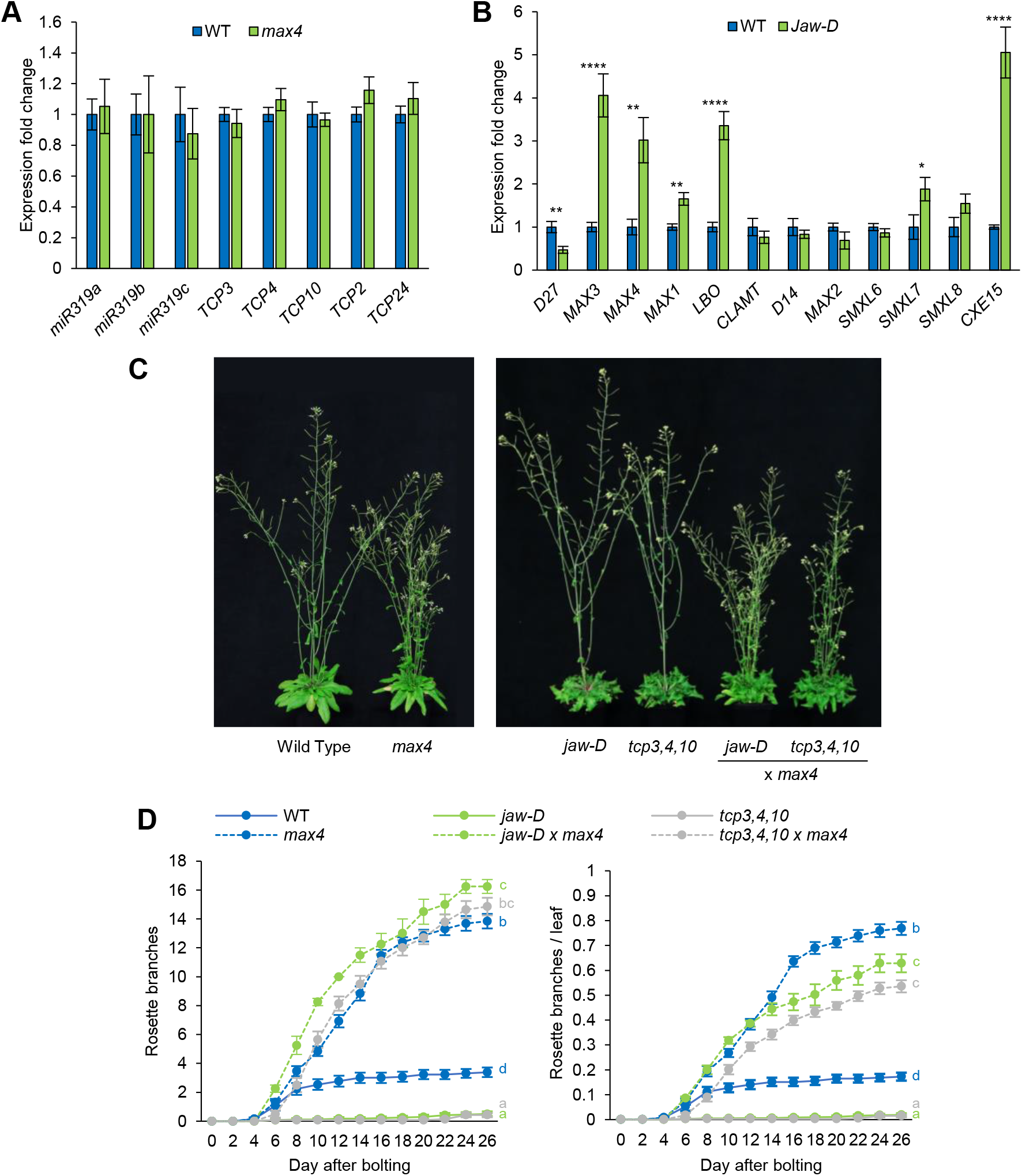
The strigolactone deficient mutant *max4* does not require TCP3, 4 and 10 to develop rosette branches in *Arabidopsis thaliana*. (**A**) expression fold change of premature *miR319* genes and their targets in three-week-old arabidopsis *max4-1* rosettes compared to the wild type. Data are mean +/-S.E. (n=8 plants). (**B**) expression fold change of strigolactone-related genes in three-week-old *jaw-D* rosette cores compared to the wild type. Data are mean +/-S.E. (n=8 plants). (**C**) shoot phenotype of the *jaw-D* and *tcp3,4,10* plants in WT Col-0 (solid line) or *max4-1* (dashed line) background, two weeks after bolting. (**D**) number of primary rosette branches (left plot) and number of primary rosette branches divided by the number of rosette leaves (right plot) emerging after bolting in *jaw-D* and *tcp3,4,10* plants crossed with *max4-1.* Data are mean +/-S.E. (n=12-20 plants). Letters indicate significant differences between genotypes at day 26 (One-Way ANOVA).

**Figure 5.**
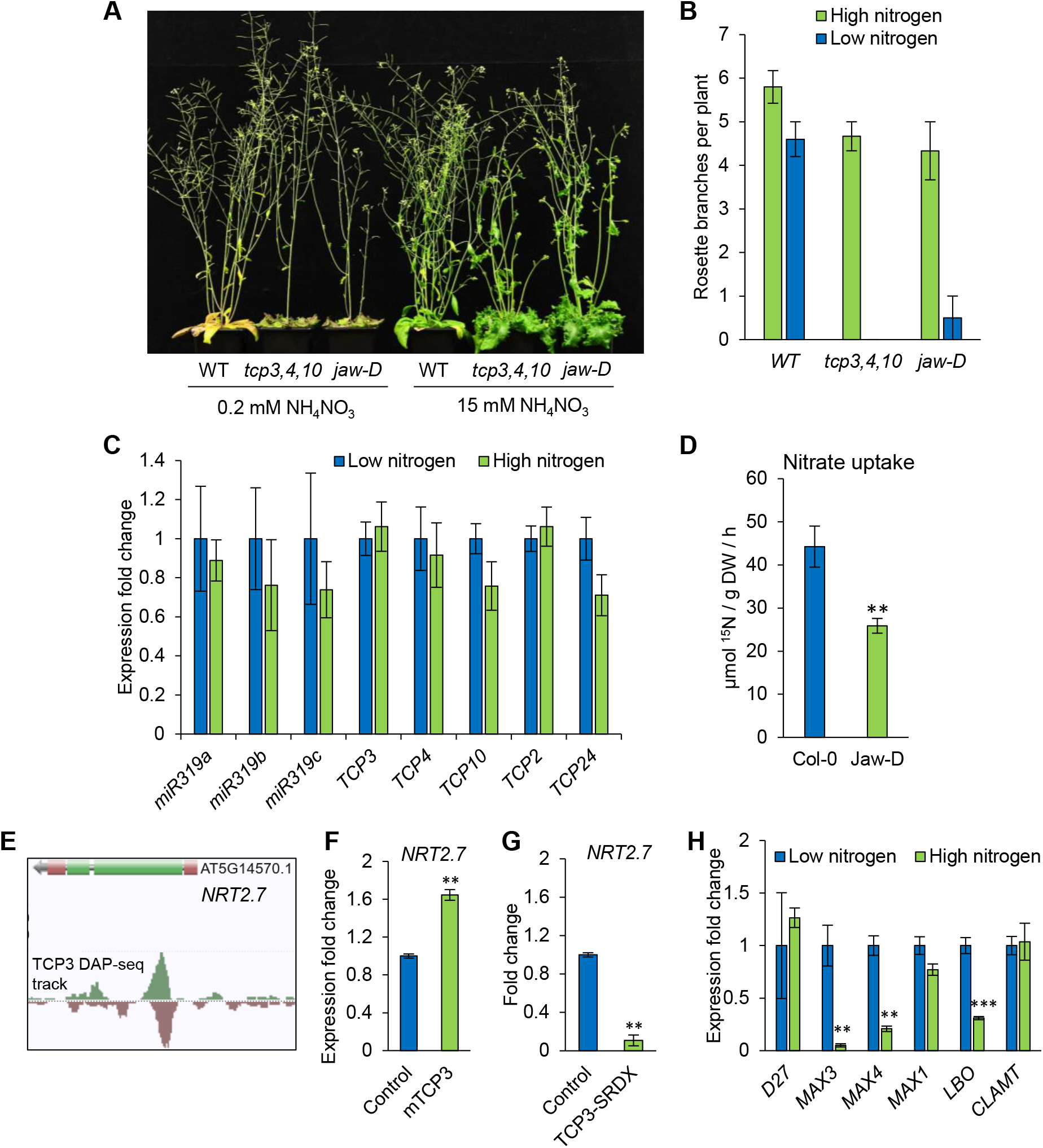
The miR319 module is involved in shoot branching response to nitrogen limitation. (**A**) shoot phenotype and (**B**) final number of primary rosette branches emerging after bolting in the wild-type (Col-0), *jaw-D* and *tcp3,4,10* KO plants supplied with either 0.2 mM (low nitrogen) or 15 mM ammonium nitrate (NH_4_NO_3_) (high nitrogen). (**C**) relative expression of *miR319* genes and their targets in arabidopsis rosettes fed with 10 mM KCl (low nitrogen) or 10 mM KNO_3_ + 10 mM NH_4_NO_3_ (high nitrogen). Data are mean +/-S.E. (n=8 plants). (**D**) nitrate root utake in 12 day old WT and *jaw-D* seedlings. Data are mean +/-S.E. (n=10-12 plants). (**E**) TCP3 DAP-seq track around *NRT2.7* genomic locus. (**F**) and (**G**) expression fold change of *NRT2.7* in plants in which *TCP3* was induced for 16 hours using a *proXVE:mTCP3* expressing line and in which TCP3 targets were repressed using a *TCP3-SRDX* expressing line, as described in Koyama et al., 2010 and 2025. (**H**) expression fold change of strigolactones synthesis genes in arabidopsis shoot supplied with high nitrogen for 60 min compared to plant supplied with low nitrogen, as described in Varala *et al*., 2018. Data are mean +/-S.E. (n=3 biological replicates).

We then determined whether the miR319 module could control shoot branching by regulating the SL pathway. To achieve this, we measured the expression of SL synthesis, signalling and degradation genes in *jaw-D* rosette cores compared to wild-type plants (Fig. 4B). The results showed that the expression of SL synthesis genes *MAX3*, *MAX4*, *MAX1* and *LBO* is significantly increased in *jaw-D*, suggesting that the miR319-target TCPs repress SL synthesis. A similar trend was observed for the SL signalling gene *SMXL7*, and the SL degradation gene *CXE15*. Results from other studies reported that the expression of these two genes is induced by SL (Wang et al., 2020; Xu et al., 2021), further suggesting that SL synthesis is increased in the *jaw-D* line.

The regulation of SL-related genes in *jaw-D* (Fig. 4A and B) suggests that the miR319 module does not act downstream of the SL pathway but may rather act, at least partly, upstream of this hormonal pathway. To clarify the relationship between the miR319 module and the SL pathway, we crossed the miR319-overexpressing line *jaw-D* and the *tcp3,4,10* triple KO mutant to the SL-deficient mutant *max4-1*. As previously reported, *max4-1* displayed a strongly increased shoot branching phenotype, with rapid release of rosette buds after bolting (Fig. 4C and D). Importantly, the *max4-1* mutation largely overrode the reduced-branching phenotypes conferred by the *miR319* overexpression and the *tcp3,4,10* triple KO mutation (Fig. 4C and D). This indicates that the positive impact of SL deficiency on shoot branching is largely independent of the miR319-targeted TCPs. Furthermore, this observation indicates that *max4* mutation is epistatic to miR319, supporting the idea that the miR319-module acts upstream of the SL pathway.

In mutants branching at their maximal capacity like *max4*, the number of rosette leaves produced strongly influences the number of primary rosette branches (Fichtner et al., 2022). We therefore normalised the shoot branching phenotype of the lines in *max4-1* background by dividing the number of rosette branches by the number of rosette leaves. Using this normalization method, we could observe a small inhibitory effect of the *jaw-D* and *tcp3,4,10* mutations on the shoot branching phenotype of *max4-1* (Fig. 4D), indicating that the miR319 module may not solely act through the SL pathway to control shoot branching.

The enhanced expression of SL biosynthesis genes (Fig. 4) is consistent with the hypothesis that SL synthesis is increased in the *jaw-D* line and that the miR319 module may act, at least partly, upstream of this hormonal pathway. However, high SL gene expression may also occur due to feedback regulation of the pathway under low SL levels (Johnson et al., 2006; Hayward et al., 2009; Waters et al., 2012). To investigate whether this effect of the miR319/TCP3 module on SL biosynthesis gene expression could be direct, we analysed available TCP3 DAP-seq data around the *MAX3*, *MAX4*, *MAX1* and *LBO* loci. Using this approach, we did not detect any TCP3 DAP-seq peaks on these genomic loci (Supp. Fig. S6), indicating that TCP3 does not bind to these regions, at least *in vitro*, and suggesting that the impact of the miR319 module on SL biosynthesis is unlikely to result from direct transcriptional regulation of these genes by TCP3.

### MiR319 increases the impact of nitrogen starvation on shoot branching

Our data indicate that the miR319 module affects SL synthesis (Fig. 4B) and that this effect is likely to be indirect (Supp. Fig. S6). Since SL levels in arabidopsis have been shown to be tightly regulated by nitrogen availability (Sakioka and Yoneyama, 2025), we tested whether the effect of the miR319 module on the SL pathway could be due to an effect on the plant nitrogen status. To test this, we first supplied *jaw-D* and *tcp3,4,10* mutant lines with two contrasted concentrations of ammonium nitrate (0.2 mM and 15 mM). Under the higher nitrogen condition, the inhibition of shoot branching was moderate in *jaw-D* and *tcp3,4,10* compared to the wild type (Fig. 5A-B). However, when grown under low nitrogen condition, which only moderately inhibited the final number of rosette branches produced by the WT (∼20%), this inhibition of shoot branching was almost total for *jaw-D* and *tcp3,4,10* lines. This result shows that deficiency in miR319-targeted TCPs increases the sensitivity of shoot branching to nitrogen limitation.

The previous result suggests that miR319 targets are likely to be involved in the shoot branching response to nitrogen. To further investigate this, we tested whether nitrogen could regulate the expression of *miR319* genes and its targets. To achieve this, we measured the expression of these genes in Col-0 plants cultivated hydroponically on contrasted nitrate concentrations. Gene expression analysis showed only a moderate and often non-significant reduction of the expression of *miR319* genes and its targets by the higher nitrogen concentration (Fig. 5C), indicating that these genes are not transcriptionally regulated by nitrogen.

We then tested whether the increased sensitivity of *jaw-D* to nitrogen limitation could be due to a deficiency in nitrate root uptake. To test this, we measured nitrate root uptake in this line, using ^15^N-lablelled nitrate at a low concentration that enables to test the high affinity transport of this nutrient (0.2 mM), as the strongest inhibition of shoot branching was observed under low nitrogen conditions (Fig. 5A). The results of this assay show that the high affinity nitrate root uptake in *jaw-D* is significantly inhibited (∼43% decrease) compared to wild-type plants (Fig. 5D).

To investigate the possible molecular basis of this reduced nitrate uptake capacity when the miR319 targets are inhibited, we assessed whether TCP3 could directly target nitrate transporters. To do so, we screened for the presence of TCP3 DAP-seq peaks around high affinity nitrate transporter genes, including *NRT2*s and *NRT1.1* (Morales de Los Ríos et al., 2026). This approach identified *NRT2.7* as a potential target of TCP3 on the CDS of this gene (Fig. 5E and Supp. Fig. S7). Analysis of other TCP DAP-seq tracks indicate that TCP24 also binds to *NRT2.7*, indicating that several miR319-targeted TCPs are involved in this interaction. Furthermore, none of the other TCPs available on the Plant Cistrome Database displayed a peak on *NRT2.7* (Supp. Fig. S7), showing that the identified DAP-seq peaks on this gene are specific to TCP3 and TCP24.

Reanalysing previously published transcriptomic data in which *TCP3* was induced for 16 hours using a *proXVE:mTCP3* expressing line and in which TCP3 targets were repressed using a *TCP3-SRDX* expressing line (Koyama et al., 2010; Koyama et al., 2025), we observed an induction and a strong repression of *NRT2.7* expression in these lines, respectively (Fig. 5F and G). Previous studies showed that overexpression of *NRT2.7* increases nitrate root uptake while loss of this gene showed the opposite effect (Chopin et al., 2007; Armengaud et al., 2025). Together, these observations further support the idea that the loss of miR319 targets leads to impaired nitrogen uptake capacity, and suggest a role of NRT2.7 in this misregulation.

The reduced nitrate uptake capacity of *jaw-D* may provide a possible explanation of the impact of the miR319 module on the expression of SL synthesis genes. Indeed, nitrogen availability is known to repress the expression of SL genes in different species (Barbier et al., 2023). Using transcriptomic data published by Varala et al. (Varala et al., 2018), we could observe that *MAX3*, *MAX4* and *LBO* are particularly sensitive to nitrate supply in *A. thaliana* shoots (Fig. 5H), mirroring the pattern observed in *jaw-D*. We propose that the increased expression of SL synthesis genes in *jaw-D* is due to the impaired capacity of this line to uptake nitrate rather than a direct effect of miR319 targets on the transcription of these genes.

### Carbon starvation represses miR319-targeted TCPs

Sugar availability and signalling play critical roles in the control of apical dominance and shoot branching in different species including arabidopsis (Mason et al., 2014; Barbier et al., 2015a; Patil et al., 2022). We therefore wanted to investigate whether the miR319 module could be responsive to carbon availability. To do so, we correlated the expression of *TCP3* and the sugar starvation marker *ASN1* in a range a publicly available transcriptomic data in which carbon availability is modulated through different means (Fig. 6A and Supp. Table S1). This analysis revealed in a very strong and highly significant negative correlation between the expression of the sugar repressed gene *ASN1* and that of *TCP3* (Fig. 6A), indicating that *TCP3* expression correlates with carbon availability.

**Figure 6.**
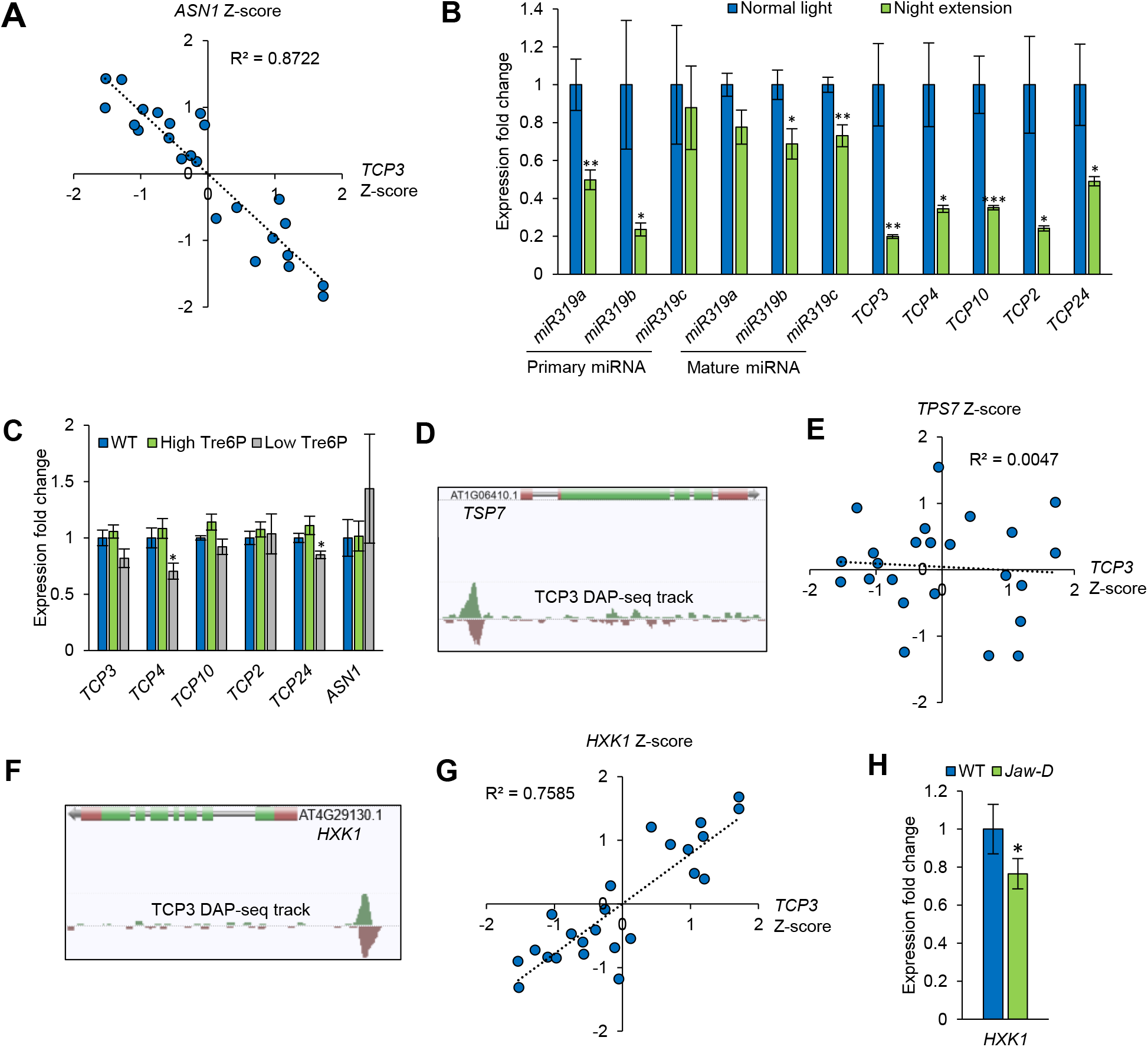
miR319 and its targets are inhibited by sugar starvation in *Arabidopsis thaliana*. (**A**) correlation between the expression of *TCP3* and the carbon starvation marker *ASPARAGNIE SYNTHASE1* (*ASN1*) in different published transcriptomic datasets in which sugar availability is modulated using various treatments (see Supp. Fig. S2). (**B**) expression fold change of the premature and mature *miR319* genes and their targets in three-week-old arabidopsis Col-0 rosettes subjected to a 6 h night extension (night extension) compared to plants that stayed in light (normal light). The expression of the premature *miRNA* and *TCP* genes was determined by RT-qPCR and the expression of the mature miRNA was obtained through small RNA sequencing. Data are mean +/-S.E. (n=6-8 plants). (**C**) expression fold change of miR319-targeted *TCPs* and *ASN1* in lines with high Tre6P (*proGLDPA:otsA*) and low Tre6P (*proGLDPA:CeTPP*) lines compared to the wild type. (**D**) TCP3 DAP-seq track around *TPS7* genomic locus. (**E**) correlation between the expression of *TCP3* and *TPS7* in different published transcriptomic datasets in which sugar availability is modulated using various treatments (see Supp. Table. S2). (**F**) TCP3 DAP-seq track around *HEXOKINASE1* (*HXK1*) genomic locus. (**G**) correlation between the expression of *TCP3* and *HXK1* in different published transcriptomic datasets in which sugar availability is modulated using various treatments (see Supp. Table S2). (**H**) *HXK1* expression fold change in *jaw-D* rosette core compared to its corresponding wild type. Data are mean +/-S.E. (n=8 plants).

We then wanted to determine whether the repression of *TCP3* expression by carbon starvation could be due to an increase in *miR319* expression levels. To test this, we quantified by RT-qPCR the expression of premature *miR319* genes and their targets in three-week-old Col-0 rosettes exposed to sugar starvation applied through a 6-hr night extension, a treatment broadly used to deplete plants from sugars (Smith and Stitt, 2007). This analysis showed that the expression of *TCP3* and its close orthologues are all strongly inhibited by the night extension treatment. This treatment also led to an inhibition of the expression of the premature *miR319a* and *miR319b* transcripts (Fig. 6B). We then determined the expression of the three mature miR319s using small RNA-sequencing. The result indicates that the mature miR319s are also slightly inhibited by carbon starvation. Altogether, these results suggest that the strong repression of *TCP3, 4, 10, 2* and *24* expression by sugar starvation is unlikely to be mediated by miR319 in arabidopsis.

### TCP3 is independent of trehalose 6-Phosphate and induces HEXOKINASE1 expression

Trehalose 6-phosphate (Tre6P) is a sucrose-specific signal, which correlates with the levels of this disaccharide, signals its availability and modulates its concentration in sink tissues (Fichtner and Lunn, 2021). Tre6P was reported to accumulate in axillary buds in pea in response to decapitation and to promote shoot branching in arabidopsis (Fichtner et al., 2021a; Fichtner and Lunn, 2021). Given that the miR319-targeted TCPs promote shoot branching (Fig. 2 and 3) and that their expression is correlated with sugar availability (Fig. 6A and B), we wanted to test whether Tre6P could regulate their expression. To do so, we determined the expression of miR319-targeted *TCP*s in transgenic lines with high (*proGLPDA:otsA*) or low (*proGLPDA:CeTPP*) Tre6P levels in the vasculature, the main site of Tre6P synthesis, as previously published (Fichtner et al., 2021a). Compared to wild-type plants, high Tre6P levels do not significantly regulate the expression of the miR319-targeted *TCP*s, while low Tre6P only moderately but significantly inhibited the expression of *TCP4* and *TCP24* (Fig. 6C).

We then sought to investigate whether TCP3 could directly target genes involved in the Tre6P pathway. To achieve this, we looked for TCP3 DAP-seq peaks around TREHALOSE-6-PHOSPHATE SYNTHASE (TPS) and TREHALOSE-6-PHOSPHATE PHOSPHATASE (TPP) genes. This search only identified TPS7 as a potential direct target of TCP3 (Fig. 6D). However, we could not observe any correlation between the expression of *TPS7* and *TCP3* using the same datasets as used in Fig. 6A (Fig. 6E), suggesting that the expression of *TPS7* is unlikely to be directly regulated by TCP3 under fluctuating carbon availability conditions. Based on these genetic and transcriptomic data (Fig. 6C and E), we propose that TCP3 is largely independent of the trehalose 6-phosphate pathway during the carbon control of shoot branching.

Besides Tre6P, the HEXOKINASE1 (HXK1) sugar signalling pathway was shown to promote shoot branching in arabidopsis (Barbier et al., 2021). To investigate whether *HXK1* is a target of TCP3, we looked for TCP3 DAP-seq peaks around this gene and could identify a clear peak on the *HXK1* promoter (Fig. 6F). This was also the case for TCP24 but not for other tested TCPs (Supp. Fig. S7). Correlation analysis between *HXK1* and *TCP3* expression indicates a positive and significant correlation between these genes using the same datasets as used in Fig. 6A (Fig. 6G). Furthermore, RT-qPCR analysis revealed that *HXK1* expression is significantly decreased in *jaw-D* rosette cores compared with wild-type plants (Fig. 6H), suggesting that the expression of *HXK1* is positively regulated by miR319-targeted TCPs.

### MiR319 over-expression partially inhibits shoot branching in the brc1 KO background

Our results indicate that the expression of *TCP3* and its close homologues are positively regulated by sugar availability (Fig. 6). In contrast, the expression of *BRC1* is strongly reduced by sugar availability, as shown by its expression in response to a 6 hr night extension (Fig. 7A), in line with previous studies reporting the same trend in different dicot and monocot species (Kebrom et al., 2012; Mason et al., 2014; Barbier et al., 2015a; González-Grandío et al., 2017; Otori et al., 2019; Patil et al., 2022). This suggests that carbon availability controls shoot branching by inducing *TCP3* and repressing *BRC1* expression. To investigate the crosstalk between these class II TCPs, we crossed the miR319-overexpressing line *jaw-D* to the *brc1* mutant. Quantification of the final number of primary rosette branches indicates that overexpression of *miR319* partly repressed the increased shoot branching phenotype of the *brc1* KO mutant (Fig. 7B and C). This result supports the idea that the balance between *TCP3* and *BRC1* expression is important for the regulation of shoot branching and that these pathways are largely independent.

**Figure 7.**
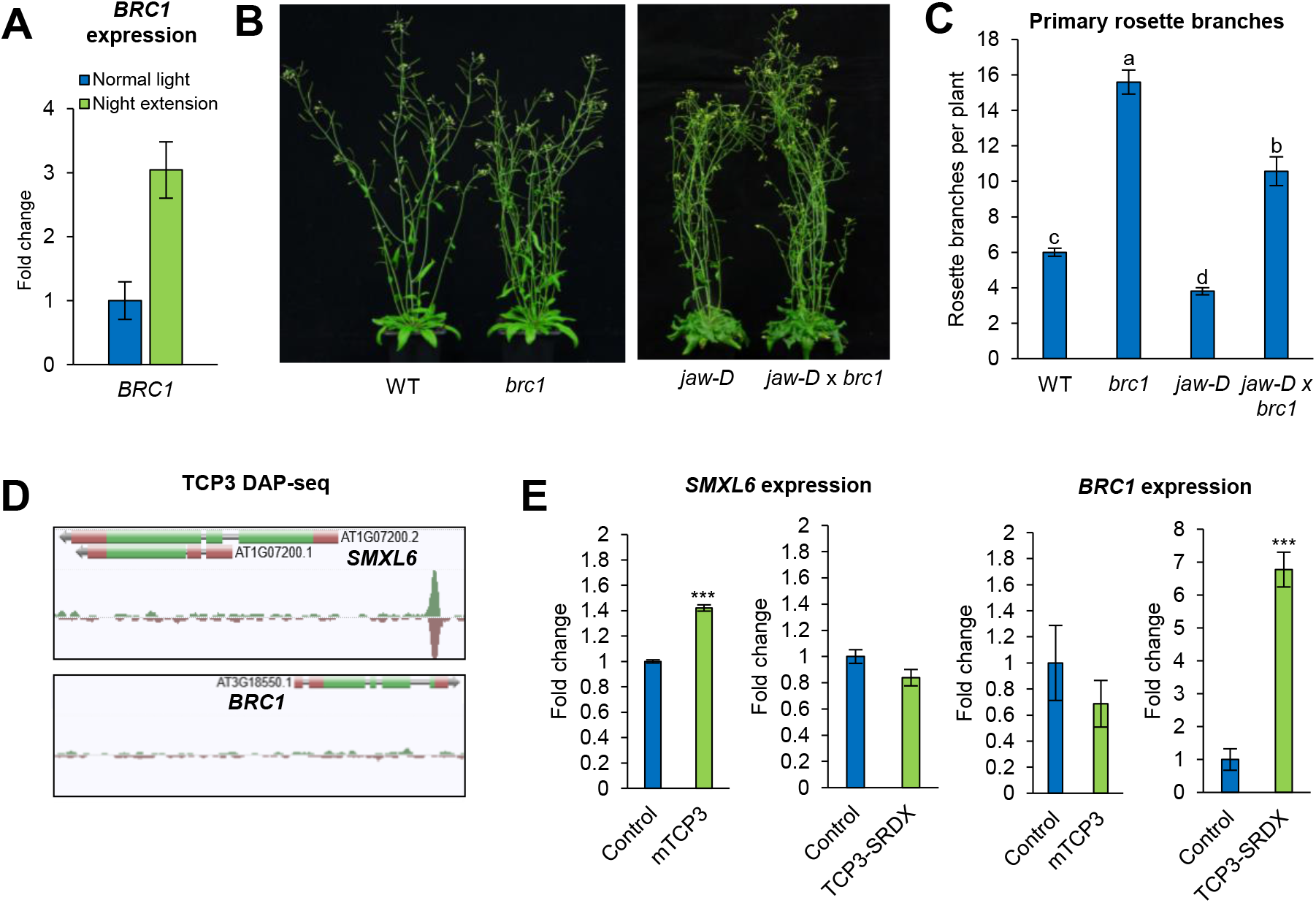
*MiR319* overexpression partly inhibits shoot branching in *brc1* KO mutant. (**A**) *BRC1* expression fold change in three-week-old arabidopsis Col-0 rosettes subjected to a 6 h night extension (night extension) or not (normal light). Data are mean +/-S.E (n=6-8 plants). (**B**) shoot phenotype and (**C**) final number of primary rosette branches of *jaw-D* plants crossed with *brc1-CRISPR* mutant. The picture was taken ∼two weeks after bolting. Data are mean +/-S.E. (n=5-15 plants). Letters indicate significant differences between genotypes (One-Way ANOVA). (**D**) TCP3 DAP-seq tracks around *SMXL6* and *BRC1* loci. (**E**) *SMXL6* and *BRC1* expression change fold in plants in which *TCP3* was induced for 16 hours using a *proXVE:mTCP3* expressing line and in which TCP3 targets were repressed using a *TCP3-SRDX* expressing line, as described in Koyama *et al*., 2010 and 2025.

A previous study reported that TCP4 binds to the *BRC1* promoter and interacts with SXML6 to inhibit the expression of this gene (Huang et al., 2026). To test whether this interaction also exists between TCP3 and *BRC1*, we looked for TCP3 DAP-seq peaks around *BRC1* and *SMXL6* genomic loci. We did not observe any DAP-seq peak around *BRC1* locus but detected one on the *SMXL6* promoter (Fig. 7D). This suggests that, unlike TCP4, TCP3 is unlikely to directly regulate *BRC1* expression, but that an inhibition of the expression of this gene remains possible via *SXML6*. This hypothesis is supported by the increased expression of *SXML6* upon TCP3 induction (Fig. 7E). Consistent with this, we observed an increased expression of *BRC1* upon induction of a repressing version of TCP3 (TCP3-SRDX) (Fig. 7E), which cannot be explained by the direct binding of TCP3 to the *BRC1* promoter, but can be explained by the binding of TCP3 to an upstream signal that inhibits the expression of *BRC1*, such as SMXL6.

## Discussion

### The miR319 module as a key regulator of primary shoot architecture

MiR319 is best known for its roles in the control of leaf morphogenesis, organ growth, floral transition, reproductive development, cell proliferation and expansion, and stress response (Palatnik et al., 2003; Nag et al., 2009; Rubio-Somoza and Weigel, 2013; Koyama et al., 2017). The results presented in this study add to the growing body of evidence that miR319 is an important regulator of primary shoot architecture in both monocotyledonous and dicotyledonous species. In rice, bentgrass, and switchgrass, *miR319* overexpression represses tillering (Zhou et al., 2013; Liu et al., 2020; Wang et al., 2021). In rice, inhibition of miR319 leads to increased tillering (Wang et al., 2021), while in wheat, it leads to the opposite phenotype (Jian et al., 2022). This difference could be attributed to miR319 monocot variants targeting distinct sets of genes. Indeed, the miR319 sequence is very similar to the one of miR159, which targets specific MYB transcription factors. Several studies in monocot species have shown that miR319 can target both TCP and MYB transcription factors (Wang et al., 2021; Jian et al., 2022). In dicots, direct evidence for the involvement of miR319 in shoot branching was lacking. Our data clearly demonstrate that miR319 inhibits axillary bud outgrowth and shoot branching in *A. thaliana* (Fig. 1 and 3), indicating that the role of miR319 in shoot branching is not limited to monocots but also extends to dicot plants. Thus, miR319 appears to be a broadly conserved regulator of plant architecture in flowering plants, although its developmental impact can differ between species.

The expression pattern of *miR319* genes appears to differ substantially between species. In rice, promoter analyses indicate that MIR319a activity is not strongly concentrated in tiller buds, but is instead detected in leaves and basal tissues, including the leaf sheath, leaf blade, and vasculature. In our study, *miR319b* and *miR319c* expression was specifically detected in arabidopsis axillary buds (Fig. 1A), revealing regulatory differences between monocots and dicots. However, in *Brassica juncea*, *miR319* promoter activity was reported mainly in vascular tissues, suggesting that even within Brassicaceae the spatial regulation of miR319 can differ (Joshi et al., 2021). These observations are important because they suggest that, while miR319 is a conserved component controlling the establishment of shoot architecture in flowering plants, its regulation and mode of action can vary greatly from one species to another. One explanation for this species-specific flexibility could be that the mobility of miR319s differ from a species to another. Another explanation could be that the miR319 module contributes to the adapted diversification of plant species to specific habitats and lifestyles. In line with this, a study in *Brassica juncea* reported that the six different miR319-encoding genes present in this species are regulated by different environmental and endogenous stimuli, indicating that the duplication and neofunctionalization of the *miR319* genes enables plants to improve their range of possible responses to the environment in order to fine-tune their development (Joshi et al., 2021).

### Class II TCP transcription factors play different roles in the control of shoot branching

The discovery of TB1 in maize and of its arabidopsis homologue BRC1 established class II TCP transcription factors as central regulators of shoot branching (Doebley et al., 1997; Aguilar-Martínez et al., 2007). In this framework, class II TCPs have largely been associated with the repression of bud outgrowth. Our results add to emerging evidence that this view is incomplete. While BRC1/TB1 represses branching, the miR319-targeted class II TCPs, and particularly TCP3, TCP4 and TCP10, act as positive regulators of shoot branching in arabidopsis, as supported by the multiple lines and genetic constructs used in this study (Fig. 1-3). Therefore, class II TCPs do not constitute a homogeneous group of branching repressors. Instead, the shoot branching network appears to rely on antagonistic activities of different class II TCPs, with BRC1 acting as a dormancy-promoting factor and miR319-targeted TCPs acting as branching-promoting factors.

MiR319-targeted TCPs have redundant functions in arabidopsis and single mutants usually have almost no phenotype. Using the triple *tcp3,4,10* KO arabidopsis mutant, we showed that this subclade of miR319-targeted TCPs is required for shoot branching, which is consistent with what was observed in a recent study (Huang et al., 2026). However, these three TCPs may control shoot branching differently. Indeed, while overexpression of a miR319-resistant version of *TCP4* did not increase shoot branching in this last study, we show that overexpression of a miR319-resistant version of *TCP3* could trigger a strong shoot branching phenotype (Fig. 2 and Supp. Fig. S3). In addition, expression of a dominant repressor version of *TCP3* (*TCP-SRDX*) driven by a constitutive *35S* promoter or by a bud-specific promoter (*proBRC1 and proSTM*) led to shoot branching repression. These observations indicate that TCP3 plays an important role in the control of shoot branching, and that this role may differ from the role played by its closest homologue TCP4.

In a recent study, Huang et al. reported that TCP4 directly binds to *BRC1* promoter to repress its transcription (Huang et al., 2026), which raises the question whether TCP3 could also act in a similar way. Our observations made with the TCP3-SRDX constructs suggest otherwise. Indeed, if TCP3 were to bind to *BRC1* promoter, overexpressing the TCP3-SRDX construct would lead to an inhibition of *BRC1* expression, thereby increasing shoot branching, which is the opposite to what we observed (Fig. 2A-B and Fig. 3A-C). Furthermore, the increased expression of *BRC1* upon TCP3-SRDX induction (Fig. 7E) and the extended expression pattern of *BRC1* observed in the *proBRC1:TCP3-SRDX* x *proBRC1:GUS* line (Supp. Fig. S4A) further supports the idea that, contrary to TCP4, TCP3 does not directly bind to the *BRC1* promoter. This idea is also supported by the absence of TCP3 DAP-seq peaks on *BRC1* promoter (Fig. 7D). However, we detected a TCP3 DAP-seq peak on the *SXML6* promoter (Fig. 7D), suggesting that TCP3 might directly control the expression of this gene, which is supported by the increased expression of this gene upon TCP3 induction (Fig. 7E). SMXL6 is targeted by strigolactone for degradation (Wang et al., 2015) and directly binds to the *BRC1* promoter to repress its transcription (Wang et al., 2020). The direct binding of TCP3 to *SMXL6* promoter provides a plausible explanation regarding the extended expression pattern of *BRC1* in the *proBRC1-TCP3-SRDX* line (Supp. Fig. S4A). Indeed, in this line *SMXL6* expression is inhibited by the *TCP3-SRDX* construct, thereby releasing the inhibition of *BRC1* by SXML6 and increasing its expression. Interestingly, a recent study in apple tree showed that the homologue of the class I TCPs, AtTCP14 and TCP15, binds to *D53*, the homologue of *SMXL6* in arabidopsis to induce its expression and promote bud break. In arabidopsis, TCP14 and TCP15 were reported to promote shoot branching in response to light quality (Gastaldi et al., 2024). Combined, our observations and those of other groups clearly demonstrate the pivotal roles played by TCP transcription factors from both classes in the control of axillary bud dormancy in response to environmental stimuli.

### The miR319-targeted TCPs link shoot branching to nitrogen availability

Beyond demonstrating a new role of the miR319-targeted TCPs in plant development, our study reveals an unexpected role for these transcription factors in nutrition and nutrient sensing. Shoot branching is well known to be sensitive to mineral nutrition, and nitrogen limitation generally restricts the outgrowth of axillary buds (de Jong et al., 2014; Sakioka and Yoneyama, 2025). The strong response of *jaw-D* and *tcp3,4,10* to low nitrogen demonstrates that this pathway contributes to the ability of plants to maintain shoot branching when nitrogen availability becomes limiting (Fig. 5A-B). This increased sensitivity to nitrogen limitation does not appear to result from a repression of miR319-targeted *TCP* expression under low nitrogen (Fig. 5C). Instead, our data suggest that the miR319 module may influence nitrogen acquisition or nitrogen status. Indeed, nitrate root uptake is strongly reduced in *jaw-D*. DAP-seq data and transcriptomic analyses indicate that the high affinity transporter *NRT2.7* is a target of the miR319 module, further indicating that this signalling pathway plays a role in nitrogen acquisition. Interestingly, TCP20 binds the promoter of *NRT2.1* and other nitrate-responsive genes, including *NIA1* and *NRT1.1*, and contributes to nitrate-dependent root foraging responses in arabidopsis (Guan et al., 2014). This raises the possibility that different TCP transcription factors from different classes contribute to distinct aspects of nitrate uptake.

Carbon availability is well known to promote high affinity nitrate transport, notably through the upregulation of *NRT2.1* expression, as demonstrated with dark and sucrose treatments (Lejay et al., 1999; Laugier et al., 2012). The bZIP transcription factor ELONGATED HYPOCOTYL5 (HY5) was demonstrated to mediate, at least partly, the impact of carbon availability on *NRT2.1* expression and nitrate root uptake (Chen et al., 2016). Our study provides a new transcription factor linking carbon availability and nitrate acquisition (Fig. 5 and 8B).

**Figure 8.**
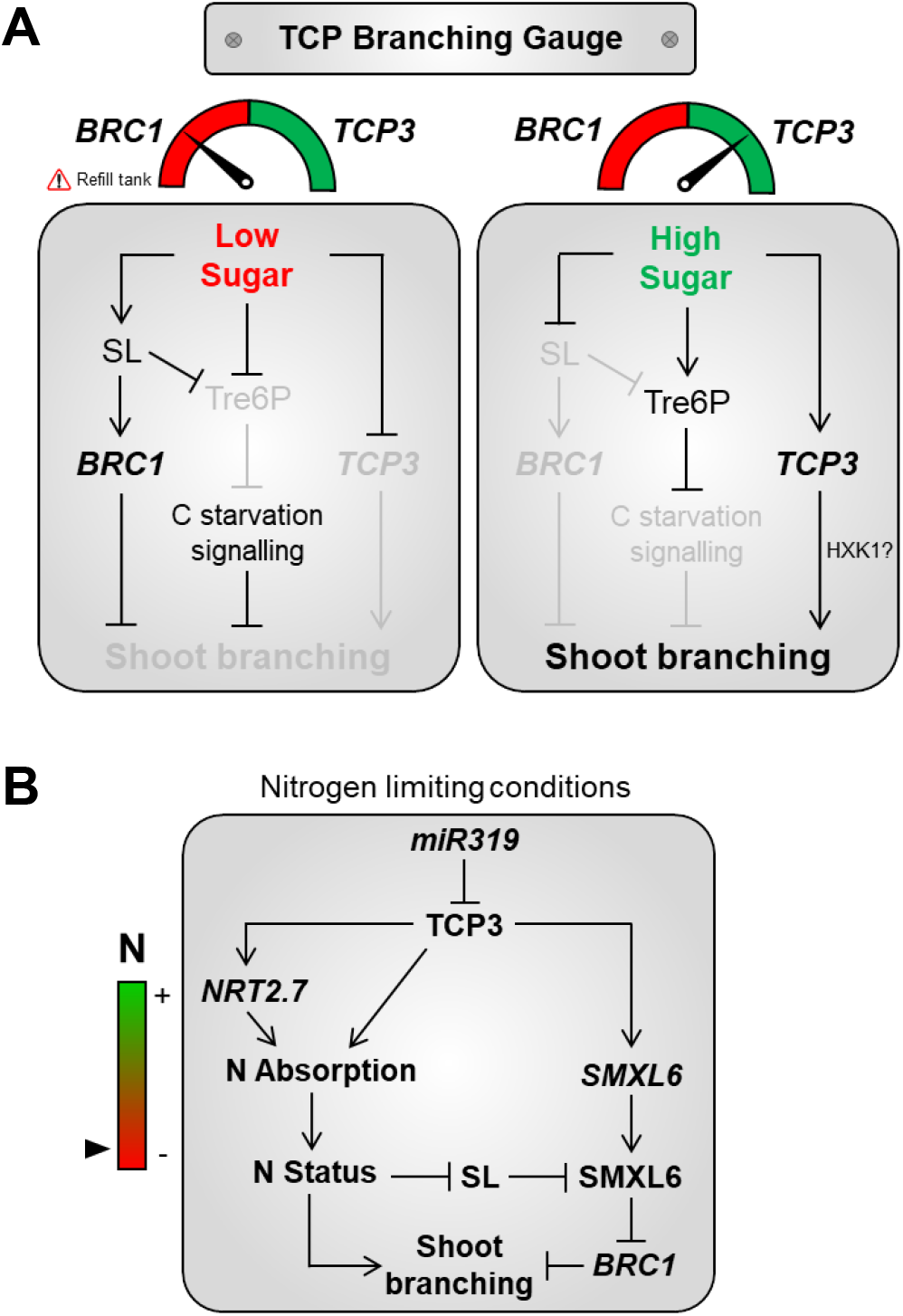
Proposed model of the involvement of TCP3 in the regulation of shoot branching in response to (A) carbon and (B) nitrogen availability.

The role of the miR319 module in nitrogen acquisition provides a plausible explanation for the observed connection between the miR319 module and strigolactone biosynthesis. In *jaw-D* rosette cores, several SL biosynthesis genes were induced, most notably *MAX3*, *MAX4*, and *LBO* (Fig. 4B). The effect of the miR319 module on these genes is unlikely to be mediated by direct transcriptional regulation by TCP3 as indicated by DAP-seq data analysis (Supp. Fig. S6). High SL gene expression can be due to low SL levels due to a negative feedback loop (Johnson et al., 2006; Hayward et al., 2009; Waters et al., 2012). If that was the case, *jaw-D* should have low levels of SLs and be bushier than the wild type, which is the opposite of what we observed (Fig. 1). Instead, we propose that the induction of SL biosynthesis genes in *jaw-D* is an indirect consequence of impaired nitrogen uptake. Indeed, reanalysis of the Varala *et al*. (2018) transcriptomic data in response to nitrogen supply showed that *MAX3*, *MAX4* and *LBO* are repressed by nitrogen supply in Arabidopsis shoots, a trend that mirrors the higher expression of these genes in *jaw-D*. Importantly, a similar relationship between nitrogen deficiency and the expression of strigolactone synthesis genes was recently reported by Sakioka and Yoneyama (2025) who showed that nitrogen deficiency influences SL levels in the basal part of shoots and affects the shoot branching phenotype in Arabidopsis. The fact that the same set of SL biosynthesis genes responds to nitrogen availability in independent datasets supports the idea that the SL-related transcriptional signature observed in *jaw-D* reflects an altered nitrogen status rather than a direct action of TCP3 on the SL biosynthetic pathway.

In this model, *miR319* overexpression reduces the activity of TCP3 and related TCPs, which compromises nitrate uptake capacity and creates a physiological state resembling nitrogen limitation (Fig. 8B). This altered nitrogen status would in turn favour the expression of SL biosynthesis genes, increasing the contribution of SLs to bud inhibition (Fig. 8B). This model is consistent with the genetic interaction between *jaw-D*, *tcp3,4,10* and *max4*. The *max4* mutation largely alleviated the branching inhibition caused by *jaw-D* and *tcp3,4,10*, indicating that SLs contribute substantially to the reduced-branching phenotype of these lines (Fig. 4C-D). However, the residual effect observed after normalising branch number by rosette leaf number suggests that the miR319 module does not act exclusively through SLs (Fig. 8). Rather, miR319-targeted TCPs likely sit upstream of several branching-related processes, including nitrogen acquisition, SL-dependent bud inhibition and possibly additional local mechanisms acting in axillary buds.

### The miR319-targeted TCPs contribute to the carbon-dependent control of shoot branching

Our results suggest that the miR319-targeted TCP module is connected to the carbon-dependent regulation of shoot branching. *TCP3* expression was positively and specifically associated with favourable carbon status across independent transcriptomic datasets and was repressed by carbon starvation (Fig. 6), contrasting with the non-responsiveness of the expression of these genes to strigolactone-deficiency (Fig. 4A) or nitrogen availability (Fig. 5C). This is consistent with the role of TCP3 and related miR319 targets as positive regulators of shoot branching, since sugar availability promotes axillary bud outgrowth in multiple species (Mason et al., 2014; Barbier et al., 2015a; Kebrom and Mullet, 2015; Otori et al., 2017; Tarancón et al., 2017). This carbon regulation is not limited to TCP3 but also affects the four other miR319-target TCPs. Surprisingly, the regulation of these TCPs by carbon availability is not simply explained by changes in premature or mature miR319 accumulation (Fig. 6B). Indeed, carbon starvation did not induce premature and mature *miR319* expression, suggesting that the decrease in TCP3 and related TCP transcripts under low-carbon conditions is more likely mediated by miR319-independent mechanisms. A previous study reported that TCP4 and miR319 act in a double negative feedback loop and that each of them is specifically expressed in distinct specific different zones of the leaf primordium (Shankar et al., 2023). It is therefore possible that our approach did not capture changes in miR319 accumulation due to this very specific spatial and temporal regulation of this miRNA and its targets. More work is needed to decipher how carbon availability controls the expression of miR319-targeted *TCP*s and to test whether this regulation is really independent of miR319.

Nonetheless, we found that the miR319 module appears to be independent of the Tre6P signalling pathway (Fig. 6C-DE), known to promote shoot branching in response to sugar availability (Fichtner et al., 2017; Fichtner et al., 2021a). This observation is important, because the relationship is reminiscent of the one reported between BRC1 and Tre6P, two signalling modules that are controlled by strigolactones and sugars but which do not act in an epistatic way (Fichtner *et al*., 2024). Our work adds TCP3 and potentially its close homologues as a new module enabling sugar availability to promote shoot branching, in parallel with BRC1-and Tre6P-dependent pathways (Fig. 7 and 8A).

Our results indicate that, contrary to the relationship between TCP3 and Tre6P, this transcription factor may interact with the HXK1-dependent pathway, also reported to promote shoot branching in response to carbon availability (Barbier et al., 2021). Indeed, our data suggest that TCP3 is likely to directly upregulate *HXK1* expression in response to carbon availability (Fig. 6F and G). This conclusion is supported by the decreased expression of *HXK1* in the *jaw-D* mutant (Fig. 6H). Interestingly, TCP3 was reported to physically interact with two catalytic subunits of SnRK1 (Nietzsche et al., 2016), a kinase that plays a central role in carbon sensing and energy management (Baena-González et al., 2007; Fichtner et al., 2021b). These observations indicate that TCP3 is tightly connected to sugar sensing and provide a framework for future studies aimed at understanding the connection between miR319-taregted TCPs and sugar signalling pathways.

Finally, our results support a model in which carbon availability regulates shoot branching through the opposite regulation of two class II TCP modules, with *BRC1* expression being repressed by carbon availability and the expression of *TCP3* and its homologues being induced by it (Fig. 8A). In addition, our data (Fig. 6C-E) combined with previous observations (Fichtner et al., 2024) indicate that these two TCP modules act in parallel with the Tre6P sugar signalling pathway during this process (Fig. 8A).

## Conclusion

This study identified the miR319-targeted TCP module as a positive regulator of shoot branching in *Arabidopsis thaliana* and shows that this module helps maintain shoot branching by supporting nitrogen acquisition. This altered nitrogen status likely explains the indirect induction of strigolactone biosynthesis genes observed when the activity of this TCP module is impaired. Finally, our data place TCP3 and related miR319 targets within the carbon-dependent control of shoot branching and highlight a likely role of these transcription factors in sugar signalling. Future work will be required to unravel the mechanisms by which carbon availability controls the expression of these *TCP*s and to determine their involvement in sugar signalling and nitrogen acquisition. Defining these roles will be essential for understanding how plants integrate carbon and nitrogen availability with hormonal pathways to shape shoot architecture.

## Acknowledgments

We would like to thank Patick Laufs for fruitful discussions and Sabine Zachgo, Thomas Jack and Tomotsugu Koyama for providing seeds for this project. This work was supported by The University of Queensland Early Career Researcher grant awarded to FB, by the DAAD-PHR PROCOPE grant awarded to FB and FF (57824938/54984VB) and by the Australian Research Council Centre of Excellence for Plant Success in Nature and Agriculture led by CB (CE200100015).

## Supplemental Material

Supp. Table S1. Primers used in this study

**Supp. Table S2.**
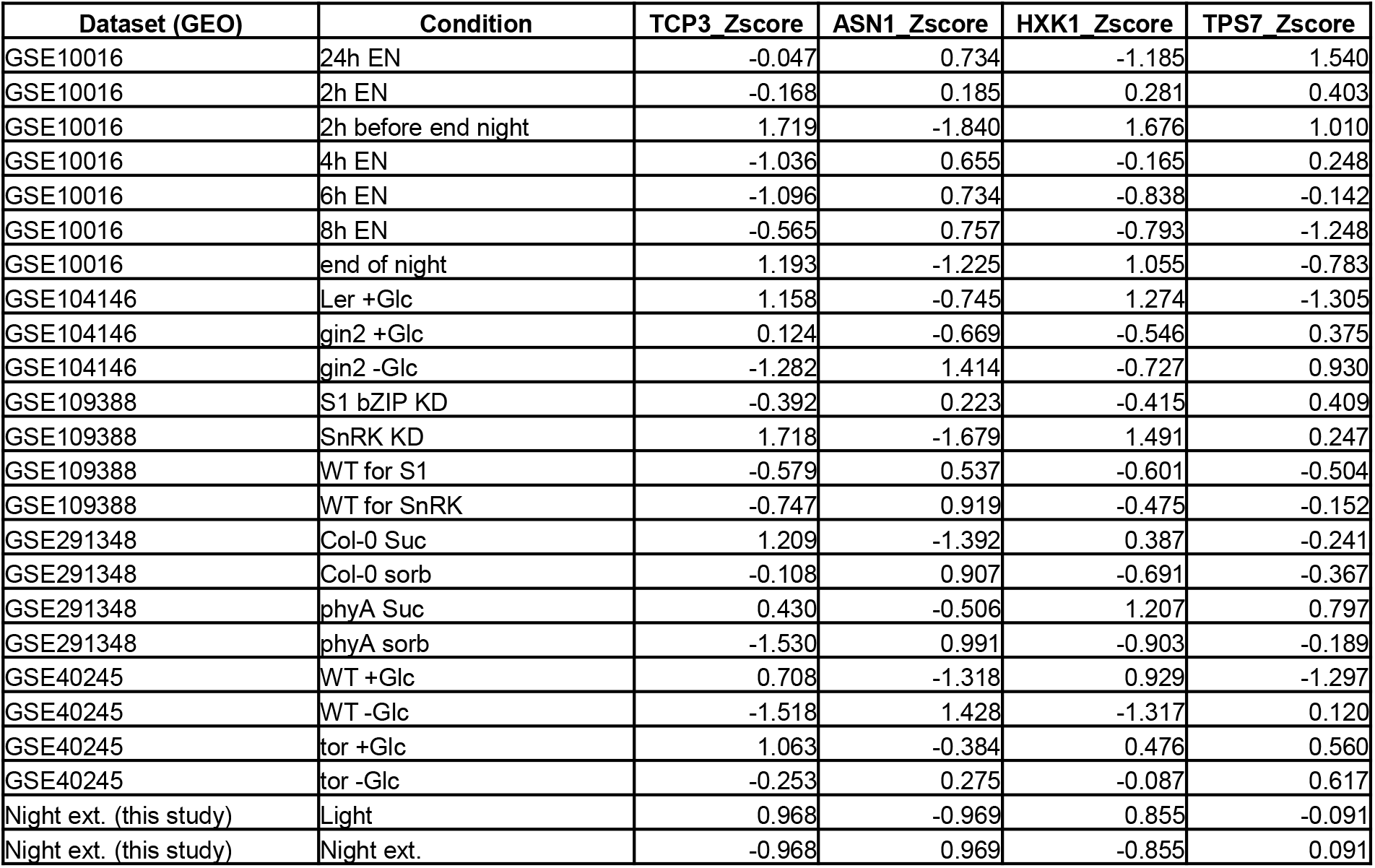
Datasets and Z-scores used in Figure 6.

| Dataset (GEO) | Condition | TCP3_Zscore | ASN1_Zscore | H XK1_Zscore | TPS7_Zscore |
| --- | --- | --- | --- | --- | --- |
| GSE10016 | 24h EN | -0.047 | 0.734 | -1.185 | 1.540 |
| GSE10016 | 2h EN | -0.168 | 0.185 | 0.281 | 0.403 |
| GSE10016 | 2h before end night | 1.719 | -1.840 | 1.676 | 1.010 |
| GSE10016 | 4h EN | -1.036 | 0.655 | -0.165 | 0.248 |
| GSE10016 | 6h EN | -1.096 | 0.734 | -0.838 | -0.142 |
| GSE10016 | 8h EN | -0.565 | 0.757 | -0.793 | -1.248 |
| GSE10016 | end of night | 1.193 | -1.225 | 1.055 | -0.783 |
| GSE104146 | Ler +Glc | 1.158 | -0.745 | 1.274 | -1.305 |
| GSE104146 | gin2 +Glc | 0.124 | -0.669 | -0.546 | 0.375 |
| GSE104146 | gin2 -Glc | -1.282 | 1.414 | -0.727 | 0.930 |
| GSE109388 | S1 bZIP KD | -0.392 | 0.223 | -0.415 | 0.409 |
| GSE109388 | SnRK KD | 1.718 | -1.679 | 1.491 | 0.247 |
| GSE109388 | WT for S1 | -0.579 | 0.537 | -0.601 | -0.504 |
| GSE109388 | WT for SnRK | -0.747 | 0.919 | -0.475 | -0.152 |
| GSE291348 | Col-0 Suc | 1.209 | -1.392 | 0.387 | -0.241 |
| GSE291348 | Col-0 sorb | -0.108 | 0.907 | -0.691 | -0.367 |
| GSE291348 | phyA Suc | 0.430 | -0.506 | 1.207 | 0.797 |
| GSE291348 | phyA sorb | -1.530 | 0.991 | -0.903 | -0.189 |
| GSE40245 | WT +Glc | 0.708 | -1.318 | 0.929 | -1.297 |
| GSE40245 | WT -Glc | -1.518 | 1.428 | -1.317 | 0.120 |
| GSE40245 | tor +Glc | 1.063 | -0.384 | 0.476 | 0.560 |
| GSE40245 | tor -Glc | -0.253 | 0.275 | -0.087 | 0.617 |
| Night ext. (this study) | Light | 0.968 | -0.969 | 0.855 | -0.091 |
| Night ext. (this study) | Night ext. | -0.968 | 0.969 | -0.855 | 0.091 |

**Supp. Figure S1.**
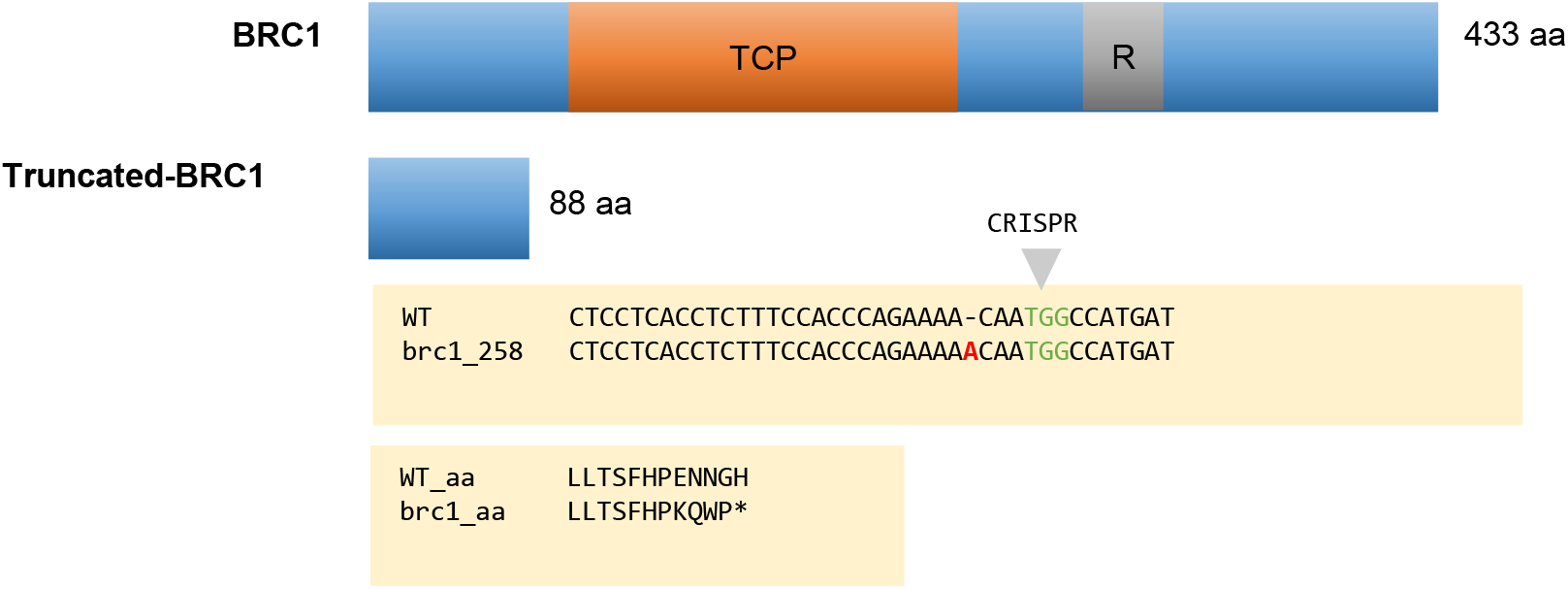
Generation of the *brc1* truncated CRISPR allele.

**Supp. Figure S2.**
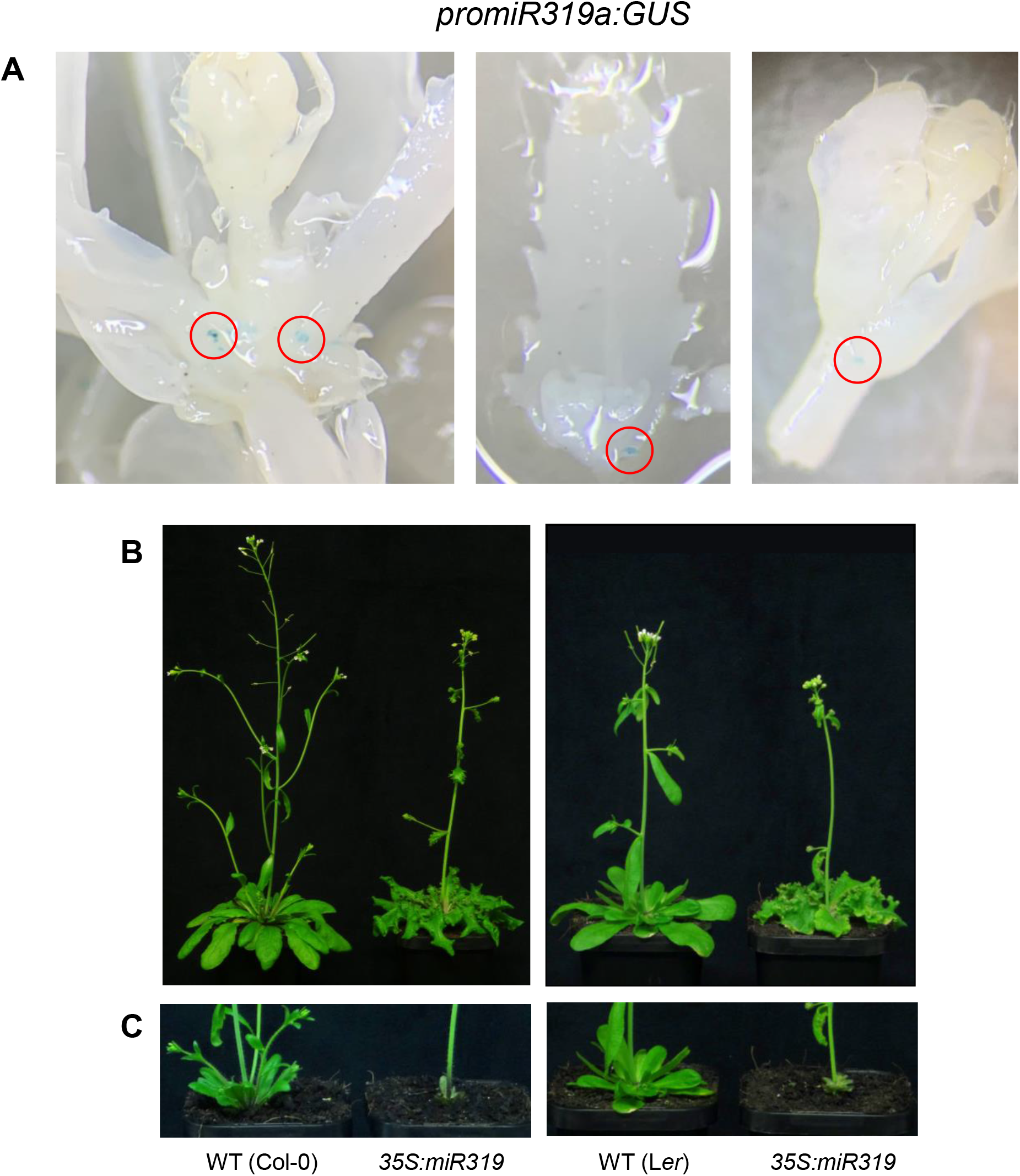
**(A**) GUS staining showing the expression pattern of *miR319a* in *Arabidopsis thaliana* rosette (**B**) shoot phenotype of (B) non-defoliated and (C) defoliated ∼6 week-old Columbia-0 (Col-0) and Landsberg *erecta* (L*er*) plants over-expressing *miR319a* compared to their respective wild-type.

**Supp. Figure S3.**
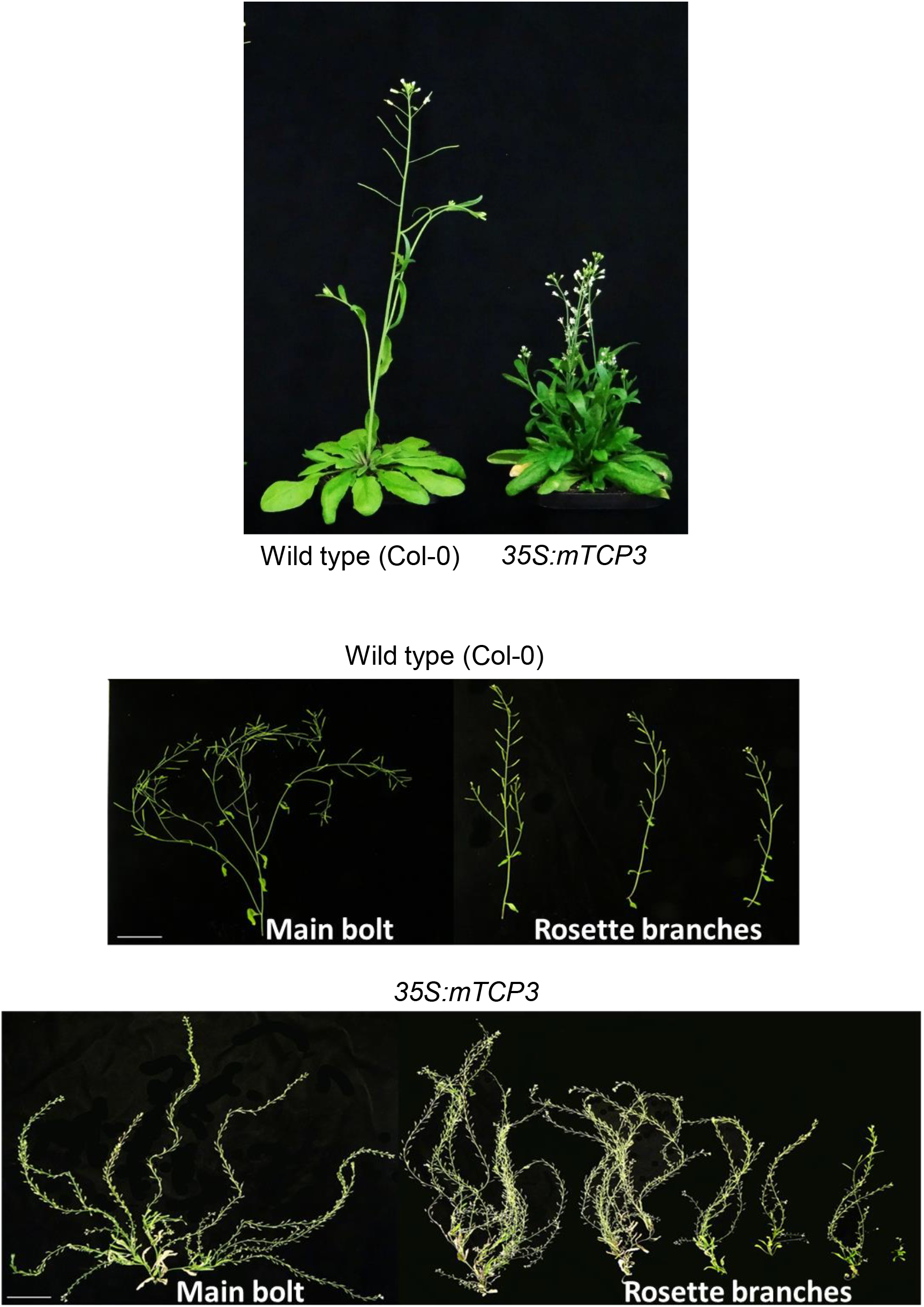
Dissection of the primary rosette branches and main bolt of the *35S:mTCP3* line.

**Supp. Figure S4.**
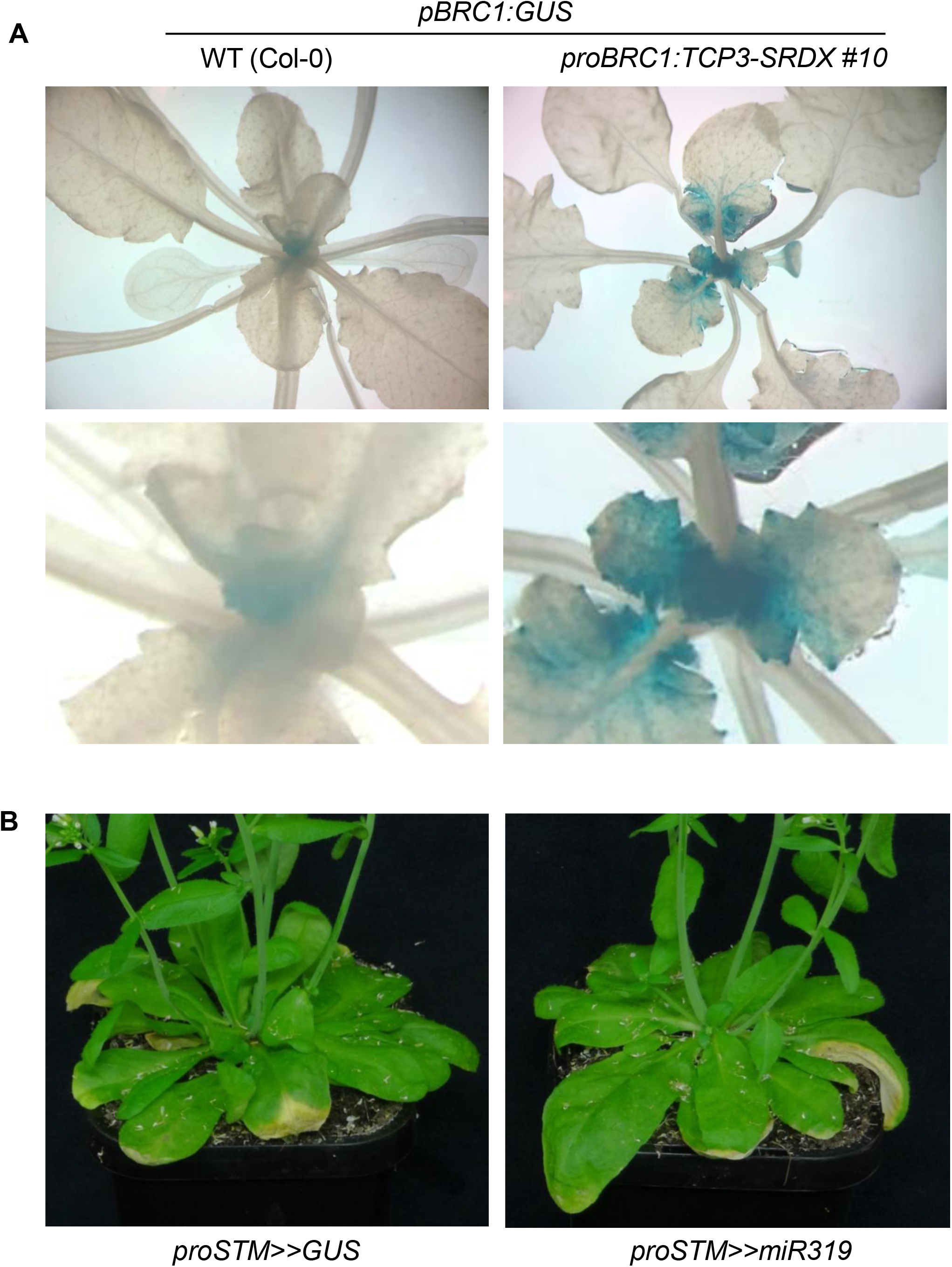
(**A**) expression patern of BRC1:GUS in Col-0 and proBRC1:TCP3-SRDX #10 background. (**B**) rosette leaf phenotype of the *proSTM>>GUS* and *proSTM>>miR319* plants displayed in Figure 3.

**Supp. Figure S5.**
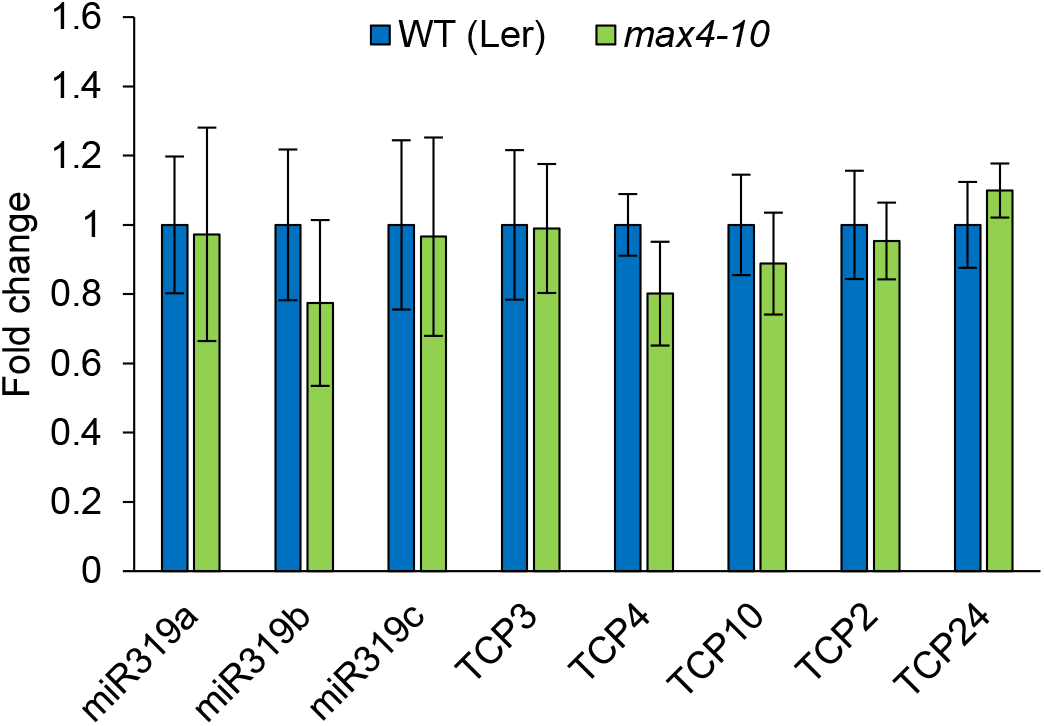
Expression of miR319 and its targets in *max-10*. expression fold change of premature *miR319* genes and their targets in three-week-old arabidopsis Landsberg *erecta* (L*er)* rosette cores. Data are mean +/-S.E. (n=8 plants).

**Supp. Fig. S6.**
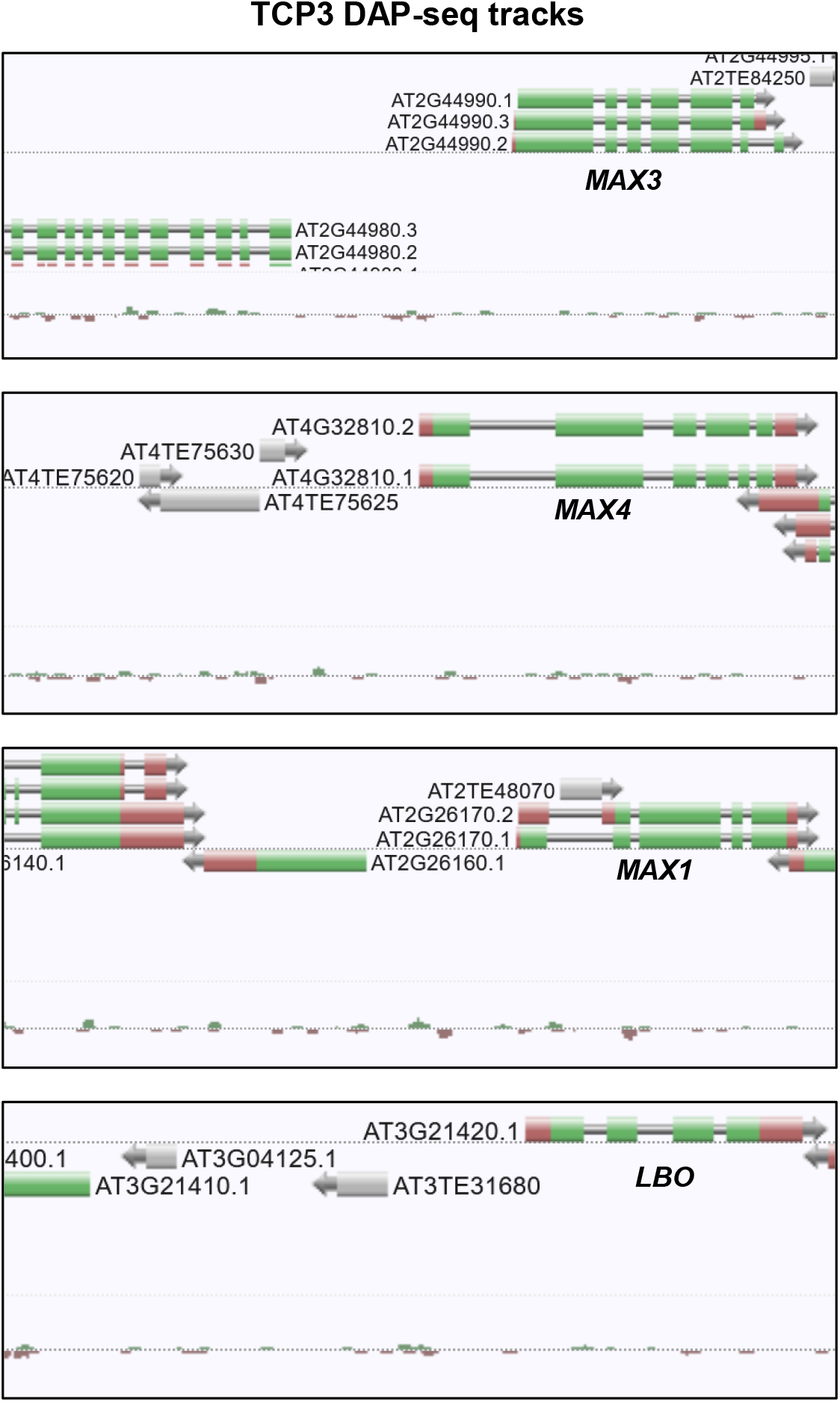
TCP3 does not bind to SL synthesis genes. Gene browser screenshot of TCP3 DAP-track around *MAX3*, *MAX4*, *MAX1* and *LBO* loci obtained

**Supp. Fig. S7.**
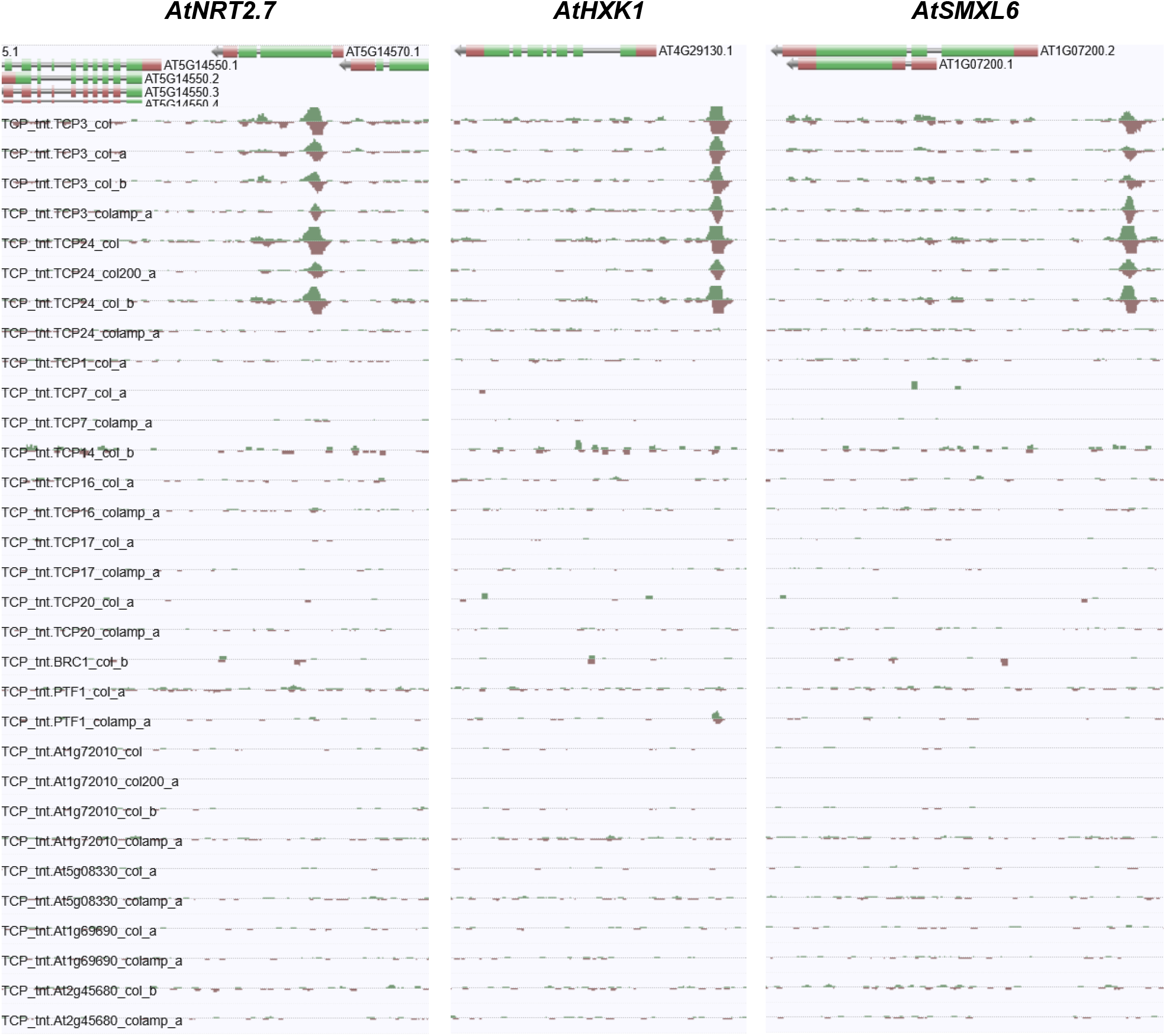
DAP-seq tracks of all the TCPs available on the Plant cistrome database around NRT2.7, HXK1 and SMXL6 loci. *O’Malley et al., 2016*

